# Genes with disrupted connectivity in the architecture of schizophrenia gene co-expression networks highlight atypical neuronal-glial interactions

**DOI:** 10.1101/2025.02.03.635993

**Authors:** Eugenia Radulescu, Petra E. Vértes, Shizhong Han, Tom M. Hyde, Joel E. Kleinman, Edward T. Bullmore, Daniel R. Weinberger

## Abstract

Dysconnectivity in schizophrenia is a pervasive trait across various levels of systems biology. To better understand disrupted patterns of molecular connectivity distinguishing schizophrenia from control non-clinical populations, we applied novel approaches to gene co-expression networks in large samples of postmortem brains from multiple regions relevant to schizophrenia: the dorso-lateral prefrontal cortex- (DLPFC) (N_donors_=297), hippocampus (N_donors_=250) and caudate (N_donors_=349). We identified differentially connected genes (DCGs) in schizophrenia networks that deviated from architectural relationships characteristic of control gene co-expression networks, by assessing three network metrics - total connectivity (K), clustering coefficient (C), and intra-module degree (kIn) determined by projecting the modular community structure of the control networks onto the schizophrenia co-expression networks. Genes showing significant absolute case-control differences for these metrics (i.e., irrespective of difference directionality) were then tested for their relationships with common genetic variants conferring risk of schizophrenia and their biological significance through post-GWAS analyses (stratified LDSC and MAGMA), gene ontology annotations and enrichment in schizophrenia-relevant gene sets.

We identified multiple DCGs, with case-control differences of connectivity metrics, consistent across brain regions. When parsed by parameter specificity, these genes show shared and specific enrichment in schizophrenia genetic signal, biological ontologies and selected cell-type markers. Notably these findings revealed widespread disturbances in co-expression connectivity affecting both neuronal and glial cells, particularly oligodendrocytes.

Overall, our results highlight disrupted co-expression network architecture in schizophrenia, implicating disrupted neuronal-glial crosstalk and its effect on synaptic transmission.

## INTRODUCTION

In E. Bleuler’s groundbreaking work “Dementia Praecox or the Group of Schizophrenias” (1) published at the turn of the 20^th^ century, he posited that schizophrenia involved a disease of “the will,” a highly complex and multi-faceted brain disorder. Decades of research since, have suggested that schizophrenia phenotypes plausibly emerge from a neurodevelopmental disorder of connectivity within the brain (2,3). Though progress has been made in unravelling schizophrenia’s biological substrate (4), much is yet to be learned about the mechanisms leading to this condition. Importantly, dysconnectivity in schizophrenia can be regarded as a pervasive trait apparent across various levels of systems biology, from genetic and molecular to cognition and behavior (5–6). The mechanisms of dysconnectivity in schizophrenia likely include transcriptional dysregulation of various genomic features (genes, non-coding regions, etc.) (7–11) that affect the optimal coordination between brain cell types and regions (12–13) and are reflected in synaptic function.

Efforts to elucidate the nature of transcriptional dysregulation associated with schizophrenia, including from our group, have focused on comparing gene expression between non-clinical (controls) and schizophrenia postmortem donor brains, and on gene co-expression network analysis (14–23). In general, these studies have been designed to find differentially expressed genes (DEG) through linear regression models or, in the case of gene co-expression studies, module detection and external validation of gene sets from modules that highlight schizophrenia risk. Because a complicated illness such as schizophrenia is a systems biology challenge, gene co-expression approaches have an appeal that differentially expressed single gene approaches cannot equal. Nevertheless, both of these approaches are limited by methodological and biological variation as discussed elsewhere (19; 24-31). A popular strategy to minimize methodological confounders and potentially disentangle the underlying biology of gene co-expression modules is to show convergence of co-expression modules across independent studies on the premise that consistent gene sets from gene co-expression modules enriched for schizophrenia genetic risk are likely more biologically relevant (20,32). This approach to date has had important, but limited success.

An alternative approach to extract the biology of risk from co-expression networks is to dig into the topological properties of the networks, to quantify the connectivity characteristics of genes within modules and within global networks and to highlight how these connectivity relationships may be altered in association with schizophrenia. In this study, we endeavored to extract intra-module, and global network metrics of connectivity within co-expression networks calculated from postmortem brain RNA-Seq, in three regions implicated in schizophrenia: dorso-lateral prefrontal cortex (DLPFC), caudate, and hippocampus. Our hypothesis is that the topological patterns of gene relationships (i.e. connectivity) in schizophrenia networks will deviate from those in control networks along specific, reproducible dimensions and these deviations will represent at a systems level the risk biology of schizophrenia. This in turn will identify novel gene sets and mechanisms potentially interesting for therapeutic development.

## RESULTS

### General characterization of control and schizophrenia co-expression networks from DLPFC, hippocampus and caudate

We created 3×2 region-by-diagnosis co-expression networks from postmortem brain bulk RNA-Seq data (brain regions: DLPFC, caudate, and hippocampus; diagnosis: controls and schizophrenia) (**Fig.1A**). In a preliminary exploratory step, we performed module detection in each network and generated six sets of co-expression modules characterized in **Fig.S1-S8** (**S1:** cluster dendrograms, number of modules per network, number of “grey” genes, **S2-S7:** biological mapping by gene ontology analysis and **S8**: module preservation- schizophrenia modules in control networks).

**Figure 1:**
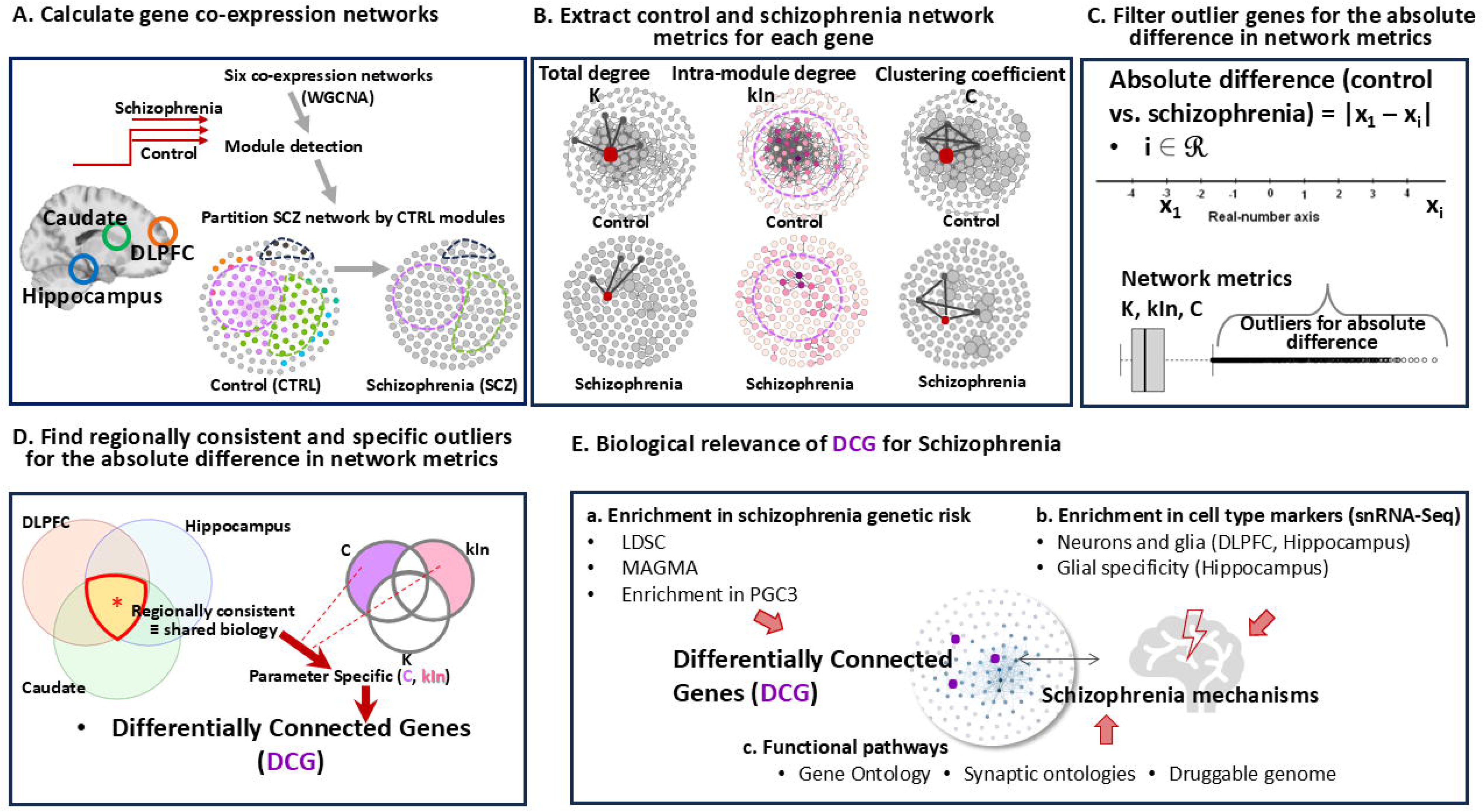
Methodological pipeline and study design. **A.** Weighted Gene Co-expression Analysis (WGCNA) on bulk RNA-Seq from postmortem brain- three regions (DLPFC, hippocampus and caudate), control and schizophrenia adult donors of European and African American ancestry; **B.** Calculation of three network parameters from each of the six networks: total degree (connectivity) K, intra-module degree (connectivity) normalized by module size (kIn), clustering coefficient (C); **C.** Calculation of each gene’s absolute difference (i.e., magnitude) in network metrics between control and schizophrenia and selecting the outlier genes by Interquartile Range (IQR) rule; outliers = genes with absolute difference above the third quartile (Q3) + 1.5*IQR, or below the first quartile (Q1) – 1.5*IQR; **D.** Assessment of regional consistency and parameter specificity of genes outliers for absolute difference in network metrics; selection of the differentially connected genes (DCG) for further functional profiling; **E.** Functional profiling and biological relevance of differentially connected genes (DCG) in relation to the risk of schizophrenia. *Legend*: DLPFC = Dorso-lateral Prefrontal Cortex; PGC3 = Psychiatric Genomic Consortium 3; snRNA-Seq = single nuclei RNA-seq; LDSC = Linkage Disequilibrium Score Regression, MAGMA = Multi-marker Analysis of GenoMic Annotation)

Biological profiling by gene ontology analysis showed similar themes for all co-expression modules, plausibly explained by the mixture of CNS cellular types characteristic for postmortem bulk RNA-seq data. Consistent with our previous studies (18–19), we observed biological processes related to neurodevelopment and CNS functionality intertwined with more general cellular functions (i.e., regulation of transcription, translation, metabolism, immunological processes) (**Fig.S2-7**).

A module preservation analysis showed that, in general, schizophrenia network modules were well preserved in control co-expression networks (Z_score_ >= 7 for all three regions) (**Fig.S8**).

### Calculation of gene-wise network metrics from control and schizophrenia co-expression networks

As illustrated in **Fig.1B**, we calculated three network metrics for each gene within each region-by-diagnosis network:

1) *Intra-modular degree (connectivity) (kIn)*. kIn is a measure that estimates how connected (co-expressed) a gene is relative to other genes in its module. The measure therefore depends on first partitioning the co-expression network into modules (see methods). While it may be interesting to compare modular partition between the CTRL and schizophrenia networks, interpreting any differences is complicated by the probabilistic nature of module detection methods and their relative sensitivity to various parameters. Here we propose to simplify the problem by using a single modular partition of the CTRL network as a reference and comparing intra-modular connectivity of all genes with respect to this same partition in both schizophrenia and control networks. We posit that this method will highlight genes whose modular affiliation is likely to change in schizophrenia, without introducing noise by running multiple module decompositions.

2). *Total degree, or connectivity (K)* is a measure of how connected a gene is relative to every other gene in the global network, irrespective of module membership. Essentially, a gene’s K total is the sum of connections strengths with all the other genes in the global network.

3) *Clustering coefficient (C)*, also independent of module membership, is a measure of the network’s local density (cliquishness) showing how interconnected a gene’s direct neighbors are.

To explore gene-specific differences in network metrics between controls and schizophrenia, we first selected as parameter the absolute difference, a measure that summarizes the magnitude of difference, irrespective of directionality (controls larger than schizophrenia or vice-versa) (**fig.1C**). In this way we generated three region specific sets of absolute difference per parameter. As shown in **Fig.2**, the absolute difference for each metric across the three regions had a skewed distribution, with most of the values close to the left tail of histograms. Correlation analysis (method Kendall tau, suitable for variables with non-Gaussian distribution) showed significant correlations between absolute differences across regions, results that indicate considerable consistency for the differences in network metrics between control and schizophrenia across the three regions (**Fig.2**, **Fig.S9-11**). We then sought to determine an appropriate threshold at which the absolute difference in network metrics becomes potentially significant for the reorganization of co-expression architecture in schizophrenia, and consequently, for disruptions in relevant biological pathways. As a threshold, we selected the upper bound defined by 1.5 times the interquartile range (IQR) above the third quartile (Q3), which is the upper boundary before individual points (i.e., genes) are considered outliers for the absolute differences in network metrics (see Methods and Fig. 1C for details).

**Figure 2:**
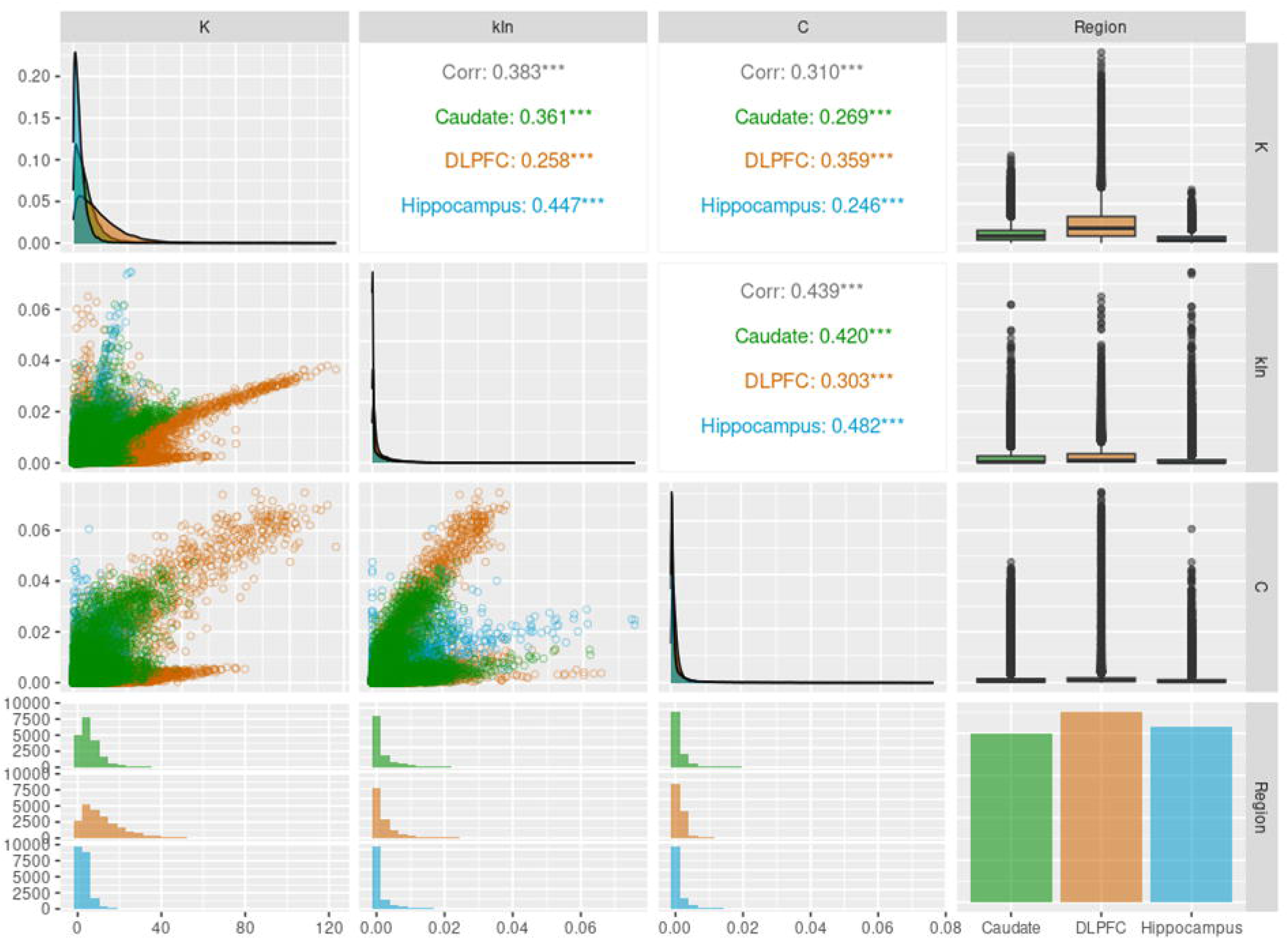
Scatter plot matrix summarizing relationships between network parameters across the three brain regions. Pair-wise correlations (method Kendall tau) (upper part of the matrix) indicate overall and intra-regional significant correlations between the three parameters; on the diagonal: density plots of parameters across regions; on the bottom axis: histograms showing the abnormal distribution of the three parameters across regions; on the right axis: boxplots showing numerous outliers for each parameter and region and a bar plot of the three regions. Scatter plot matrix created with the *ggpairs* function from the *GGally* R package (v.2.2.1).

### Identification of outlier genes for differences in network parameters in CTRL vs. schizophrenia

The next steps were to select and functionally characterize genes (nodes) with the largest differences in network parameters between CTRL and schizophrenia, specifically the outlier genes for the absolute difference in network metrics (**Fig.1C)**. As shown in **Fig.3**, numerous genes were found to be outliers for each network parameter, especially within hippocampus, followed by the caudate and DLPFC. Some of the differences in the number of outliers may be explained by the variable cell type architecture between the three regions that starts in early neurodevelopment (33) and consolidates during adulthood (34), although variation in the quality of RNA-Seq data cannot be completely ruled out.

**Figure 3:**
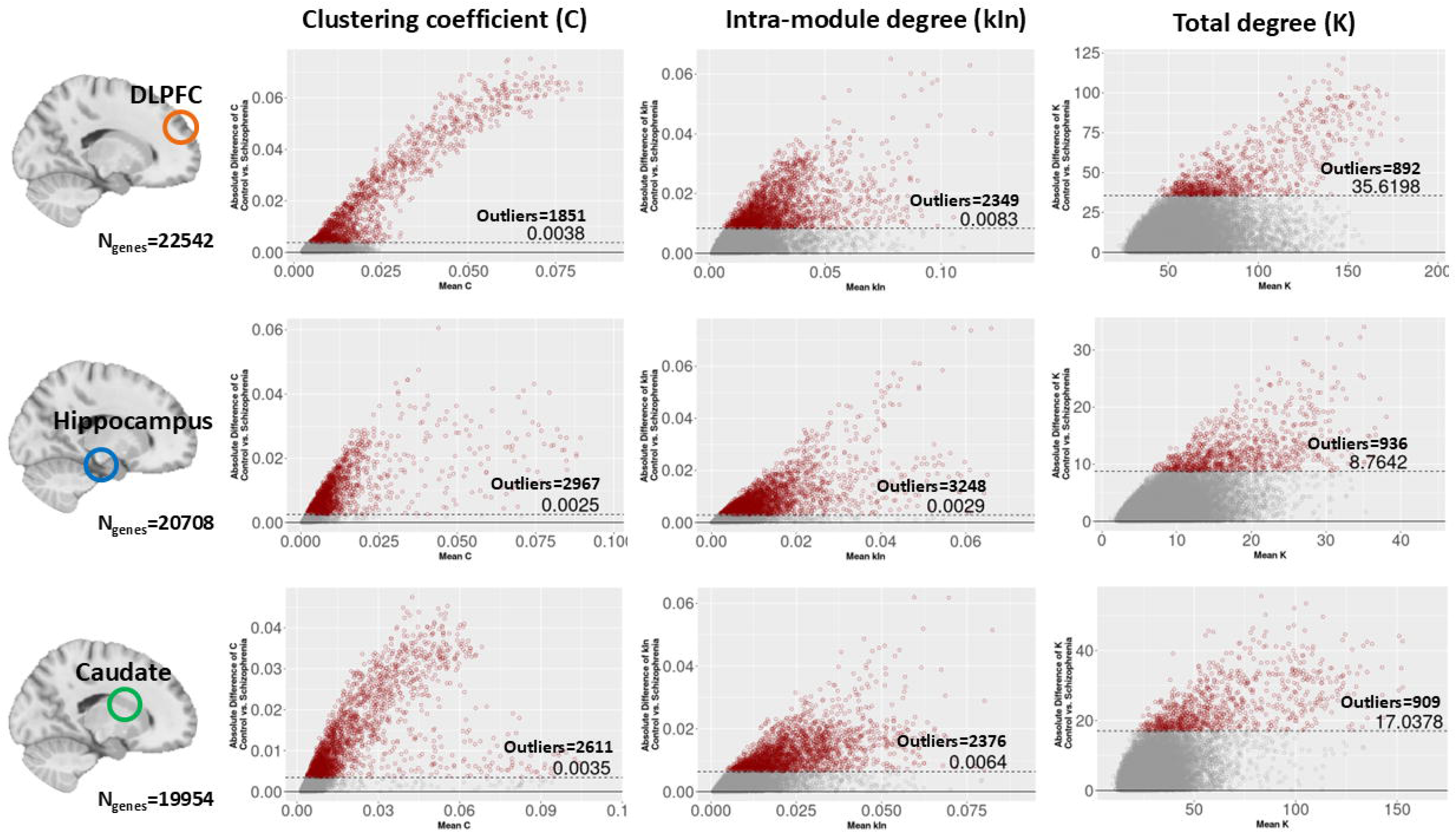
Scatter plots of case-control absolute difference in network metrics. On x-axis: the average of parameter for the two groups, control and schizophrenia; on y-axis: the absolute difference; the dashed horizontal lines mark the limits above which lie the outliers for the absolute difference calculated with IQR rule; the dark-red points represent the outliers for the absolute difference.

For functional characterization, we first filtered the outlier genes by brain regional consistency (**Fig.1D**). We reasoned that outlier genes common to all three regions potentially share a similar biology and are less likely driven by technical variation and cell composition. We then hypothesized that within the regionally consistent outliers, each network parameter may capture specific aspects of biological mechanisms underlying structural abnormalities in schizophrenia co-expression networks.

Notably, notwithstanding some variation, the three brain regions shared a sizable proportion of outlier genes, as demonstrated by the results of the fold enrichment (FE) test (**Fig.4A, 4B** and **4C**). The outlier genes common between all three regions were considered biologically credible sets and were then split by network metrics’ specificity. Overall, 100 regionally consistent outlier genes were common to all parameters, while 286 were C specific, and 185 were kIn specific (**Fig.4D**). For total connectivity (K), the vast majority of regionally consistent outliers were common between the three parameters, leaving only 9 genes specific to K. We, therefore, isolated regionally consistent outliers partitioned only by kIn and C specificity and used these differentially connected genes (DCG) as final sets for further functional profiling (**Fig.1D**).

**Figure 4:**
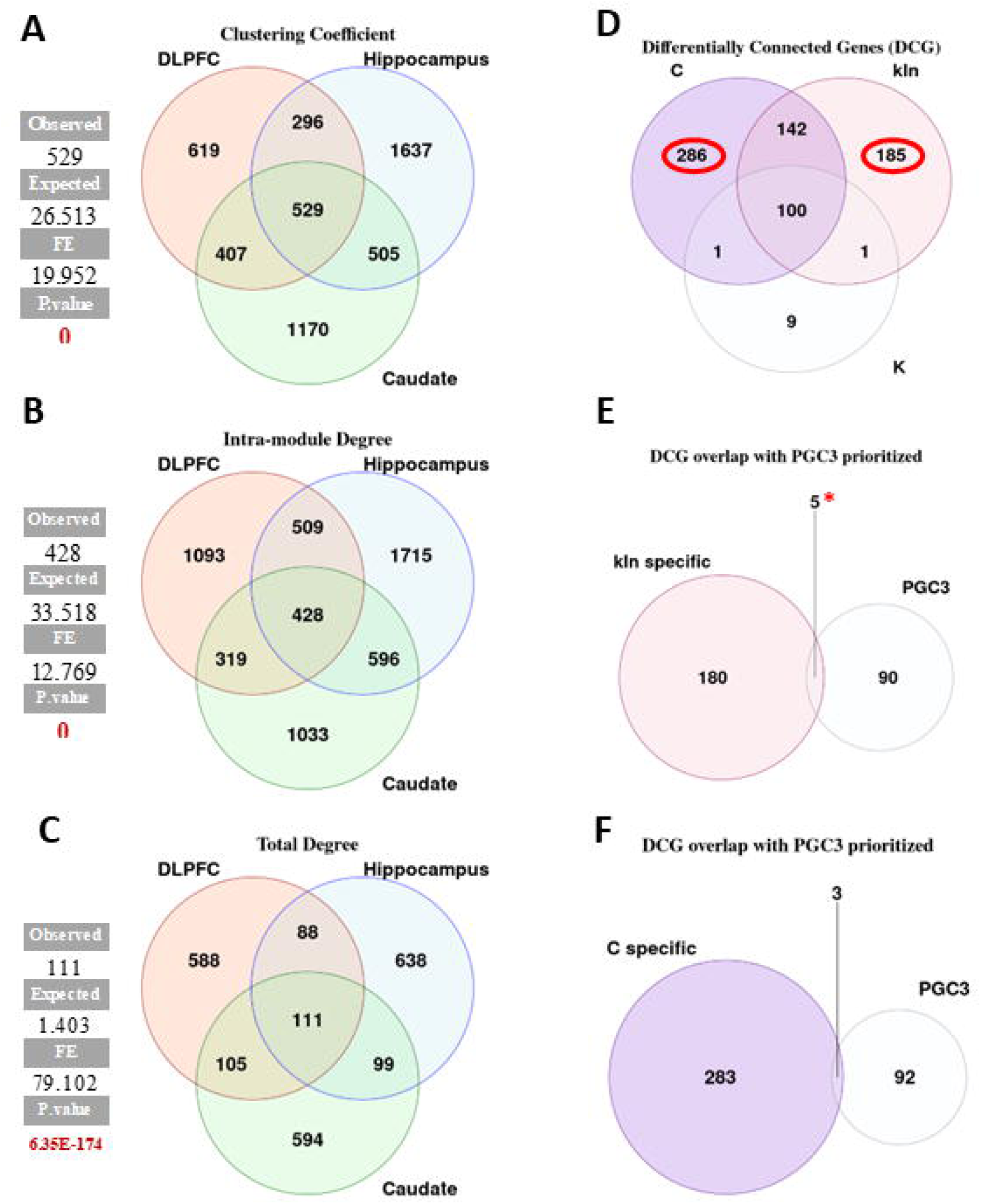
Regionally consistent and parameter specific differentially connected genes (DCG). **A-C.** Venn diagrams showing the overlap of outliers across regions; on the left- the statistical significance of overlap calculated with the fold enrichment (FE) test; D. Venn diagram showing the overlap between regionally consistent parameters (C=529, kIn=428, K=111); in red circles: the number of DCG used for further functional profiling; **E-F.** DCG overlap with the PGC3 prioritized genes (N=95 within the total of 18,874 common to the three brain regions), statistically significant for kIn DCG after FDR correction, with an overlap of five genes (**E**); C DCG overlap with PGC3 was not statistically significant after multiple comparisons correction.

### Biological profiling of schizophrenia differentially connected genes (DCG) derived from gene co-expression networks parameters

After isolating the set of differentially connected genes (DCG), we asked several questions: are these genes credibly related to biological disturbances implicated in schizophrenia? Further, do the DCG carry potential for finding schizophrenia biomarkers and drug targets?

To answer these questions, we adopted the strategy schematized in **Fig.1E**; specifically, we looked at the enrichment of DCG in schizophrenia GWAS signal, enrichment of DCG in gene sets and pathways credibly or previously linked to schizophrenia and at the overlap between DCG and cell-type specific markers determined in transcriptomic studies (single-nuclei- snRNA-Seq).

In addition, we hypothesized that DCG will bring complementary information about biological relevance for schizophrenia in comparison with more traditional gene lists derived from differential gene expression. Therefore, we selected differentially expressed genes (DEG), from previous postmortem studies of our brain regions of interest (15,35) and used them in comparison to DCG for biological profiling. This comparison also indirectly relates to the potential role of drug treatment in the DCG set, while DEGs have been shown to be dominated by drug effects in schizophrenia tissue (36).

#### Post-GWAS analysis

To look for potential relationships between DCG and mechanisms relevant for schizophrenia risk, we first checked for associations between DCG and GWAS signal for schizophrenia by applying stratified Linkage Disequilibrium Score Regression (**LDSC**) (37–38) and by gene-set analysis by Multi-marker Analysis of GenoMic Annotation (**MAGMA**) (39).

Stratified LDSC (38) is an enrichment method based on GWAS statistics of polygenic traits (i.e., schizophrenia) that estimates the enrichment of SNP-heritability for genomic annotations (i.e., in our case C and kIn differentially connected genes- DCG) based on a joint model with other annotations (i.e., coding and evolutionary conserved regions, regulatory elements and histone marks). One condition to obtain reliable results with stratified LDSC is that the gene set of interest covers at least 0.05% of SNPs of the whole genome (40). Only C-specific and kIn-specific differentially connected genes (DCG) fulfilled this condition and were therefore used for LDSC analysis.

For stratified LDSC, in addition to GWAS schizophrenia PGC3 summary stats, we also used GWAS summary statistics for a selection of other general complex traits (body-mass index- BMI, height, intelligence, smoking), and psychiatric and neurological syndromes and disorders (bipolar disorder, major depressive disorder- MDD, epilepsy, insomnia, ADHD, Alzheimer’s disorder, autism, alcohol dependence). Stratified LDSC showed enrichment in heritability of schizophrenia for both parameter specific DCG- clustering coefficient (C) (p = 0.04) and intra-modular connectivity (kIn) (p = 0.01), (**Table S1**). Notably, in addition to schizophrenia heritability, C specific DCG were significantly enriched in heritability for Alzheimer’s disease, and BMI, whereas kIn specific had a broader heritability enrichment, including bipolar disorder, alcohol dependence, epilepsy, MDD and BMI.

**MAGMA** analysis (39) is a post-GWAS functional genomics method that facilitates understanding of biological relevance of complex traits through gene-set analysis. Essentially, it models the joint GWAS effects of all the SNPs in and around a gene, considering linkage disequilibrium (LD). In this manner, genes become units of analysis and can be interrogated with respect to associated biological pathways through competitive (basic or conditional) gene-set analysis. In our study, we performed three steps of MAGMA: a) the standard gene analysis (results in **Table S2a**); b) basic competitive; c) conditional competitive. In the basic competitive gene-set analysis we used six gene sets: C and kIn specific, differentially expressed genes (**DEG**) shared by DLPFC, hippocampus and caudate, and we also added three gene-sets with synaptic ontologies (41)- SynGO genes (related to synaptic signaling, presynaptic and postsynaptic processes) based on the well documented enrichment of synaptic genes in schizophrenia genetic signal (42) that can confound a possible association between DCG and schizophrenia genetic risk. Overall, after correcting for multiple comparisons (n=6 tests), 4/6 gene-sets (kIn specific and the three SynGO sets) were significant for association with schizophrenia genetic signal (**Table S2b**). However, irrespective of statistical significance in step b), all DCG sets had genes with gene-level association evidence with schizophrenia at genome-wide significance (p<2.8 x 10^-6^): kIn specific- five genes (AKT3, SNAP91, KIAA1549, DGKZ and MAP1A), and C specific- five genes (RFTN2, C12Orf76, ARHGAP1, SPOCK3, and GRM3). The DEG set was not significant in the basic competitive Gene Set Analysis (GSA) step of MAGMA and had four genes enriched for schizophrenia signal at genome-wide significance. For conditional competitive GSA, we used our DCG of interest- C and kIn, while controlling also for SynGO genes (synaptic biological processes: pre- and postsynaptic, synaptic signaling, and synaptic metabolism). kIn specific remained significant (p = 0.018) after conditioning on each of the SynGO sets (**Table S2.c**), suggesting that kIn genetic signal is at least partially independent of some synaptic ontologies and raising the possibility of a more complex biology convergent at synaptic level.

In summary, stratified LDSC and MAGMA showed that kIn DCG were enriched for schizophrenia genetic signal by both methods, whereas C DCG set was enriched for schizophrenia genetic signal only nominal (p=0.04) by stratified LDSC. Of note, it is not unusual to have a seemingly discrepant result with these two methods, while they use different parameters to annotate genomic regions and statistical models (43).

### Enrichment of differentially connected genes (DCG) in gene sets and pathways credibly or previously linked to schizophrenia

We examined the potential biological importance of our DCG by performing gene-set enrichment analysis (GSEA) using curated gene sets, respectively: the PGC3 prioritized (i.e., functionally mapped) genes in the latest GWAS (42), genes from three TWAS studies on our brain regions of interest (15–16, 35), the sets of synaptic genes downloaded from the SynGO database (41) and gene targets for schizophrenia downloaded from the Druggable Genome database, parsed by the drug development stage (44). Results of the GSEA summarized in **Table S3a** showed that only the kIn specific genes were significantly enriched in PGC3 prioritized genes. Likewise, kIn showed significant enrichments in synaptic ontologies and druggable targets, especially for the Tbio level (i.e., target from plausible biological pathways, however, without drug or small molecules activities).

In contrast, C specific DCG did not show any significant association with our selected gene sets credibly linked to schizophrenia (**Table S3a**).

To infer a possible cell-type specificity for DCG, we used cell-type markers from two single-cell RNA-Seq datasets that cover a diversity of neuronal and glial populations: one from DLPFC and hippocampus (45) that has a more granular sub-classification of neuronal population and one from hippocampus (46) with a more in-depth characterization of glial sub-populations.

Interestingly, in this analysis, kIn specific DCG had no preferential association with a cell type population or sub-population. In contrast, C specific DCG were enriched in cell-type markers, specifically for oligodendrocytes (**Table S3b**). Likewise, C specific DCG appeared biased toward mature oligodendrocytes sub-populations, characterized in (46) as related to myelination processes and oligodendrocytes differentiation (**table S3c**).

#### Biological mapping by gene ontology (GO) analysis

GO analysis of DCG suggested a complex interaction between mechanisms implicated in the neurodevelopment and optimal functionality of CNS synapses (**Fig.5**). Mirroring the enrichment in schizophrenia related gene sets and cell-type markers, these results highlighted not just the involvement of neuronal mechanisms, but also a contribution of glial cells, especially oligodendroglia (**Fig.5**). Likewise, a relative functional divergence was observed between the DCG, with C-specific genes more enriched in biological processes related to oligodendrocytes’ development and axon ensheathment in CNS, and kIn-specific genes preferentially enriched in processes related to modulation of chemical transmission and immune functions (**Fig.5**).

**Figure 5:**
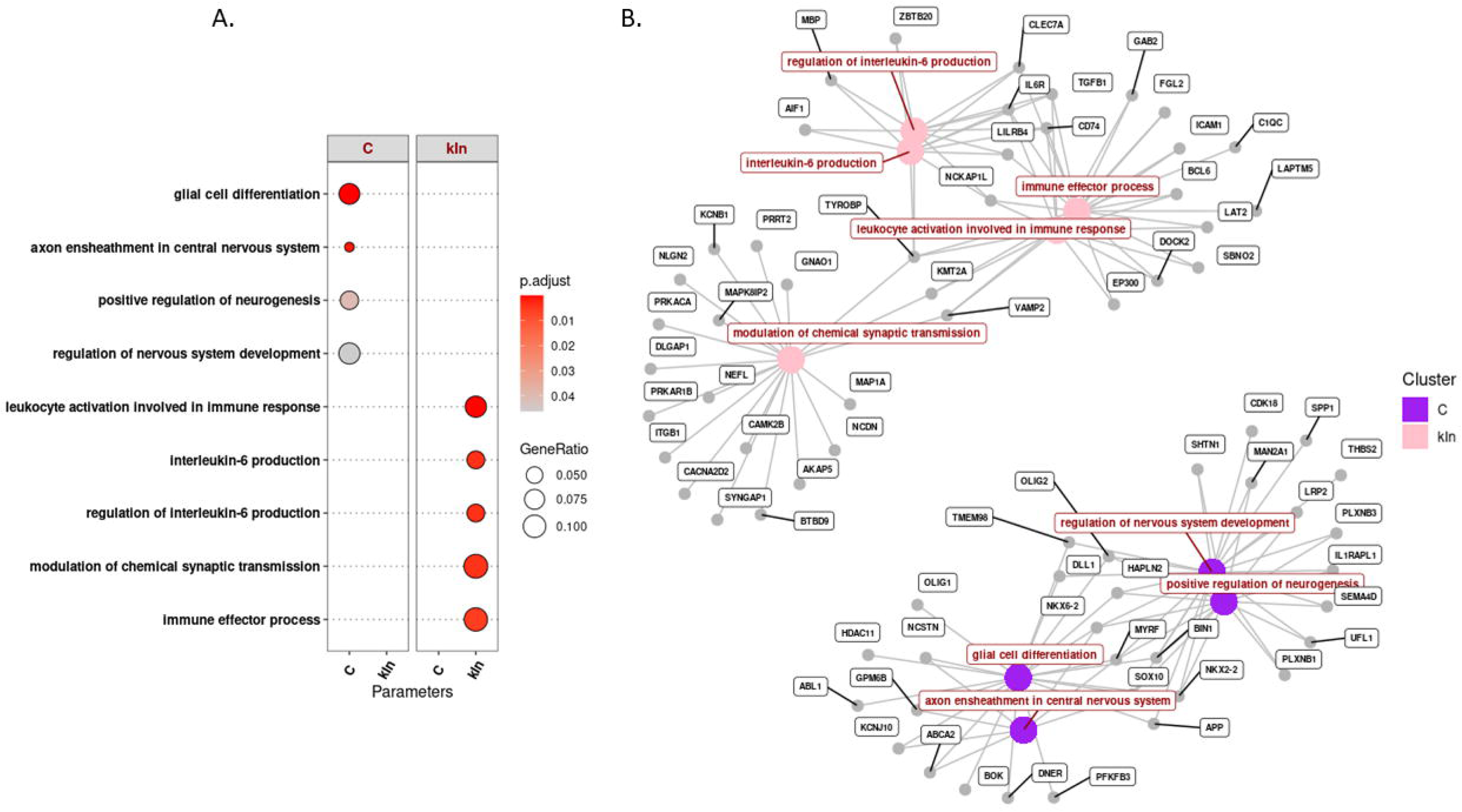
Biological annotation of differentially connected genes (DCG) by gene ontology analysis; **A.** pathway divergence between parameter specific deviant genes by gene ontology enrichment comparison; **B.** Cnetplot showing genes within biological pathways and relationships between biological ontologies (analysis and visualization with *clusterProfiler* R package).

By comparison with DCG, the DEG set had enrichment only for the druggable genome (T bio set) (**table S3a**). Likewise, DEG had no GO:BP enrichment. Importantly, there was no significant overlap between DCG and DEG. This suggests that our schizophrenia DCG derived from co-expression network metrics have the potential to bring additional information about the biological disturbances underlying schizophrenia.

## MATERIALS AND METHODS

### Data selection and processing

Postmortem human brain tissue from the LIBD Human Brain Repository was used in this study; the tissue was collected under a standardized protocol for brain acquisition, processing, and curation (location, legal authorizations, informed consent, clinical review/ diagnosis) (15–16, 35), (details in **Supplementary Methods- SM1**).

Tissue from three brain regions was analyzed: DLPFC (N_donors_=297; N_genes_=22,542), Caudate (N_donorss_=349; N_genes_=19,954) and Hippocampus (N_donors_=250; N_genes_=20,708). Donors and gene overlap for the three regions was N_donors_=155, respectively N_genes_=18,874. Data was collected from adult donors, neurotypical (control) and schizophrenia, European (Eur) and African American (AA) ancestry (**Table S4**).

RNA-seq processing consisted of quality check (QC) of each region’s dataset (i.e., removal of gene expression outlier samples), and “cleaning” (adjustment through linear models to preserve the diagnosis effects and remove unwanted variance associated with biological variables of non-interest, (i.e., demographic variables (age, sex), ancestry (10 genomic principal components) and technical artifacts (RIN, mito-rate, rRNA rate, total assigned genes, quality surrogate variables- qSVs) (18–19). Multicollinearity analysis showed that several variables were highly correlated (colinear) and were removed from the linear models based on the variation inflation factor (VIF) (**SM2**).

Linear regression models (**Eq.1**) were used for gene expression adjustment (“cleaning”), with gene expression as dependent variable and diagnosis and variables of non-interest as independent regressors. This step was performed with *cleaningY* function from the jaffelab package (47) (**SM2**).

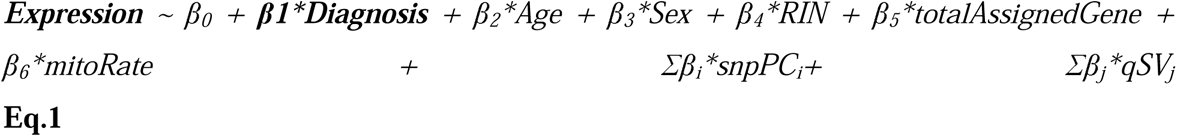

Residuals calculated from the adjustment linear regression models were used as input to create region and diagnosis specific gene co-expression networks with Weighted Gene Co-expression Network Analysis (WGCNA) (48) (**Fig.1A**).

### Creating control and schizophrenia co-expression network with WGCNA

A scale free topology analysis with 30 soft thresholding beta (β) powers (*pickSoftThreshold* function from WGCNA) was applied to control and schizophrenia expression sets to select a common beta (β) power, one for each diagnostic group. Prior to robust WGCNA, a bootstrapping procedure was performed to generate 50 controls, 50 schizophrenia random samples of gene expression by using as input residuals calculated in the expression data cleaning step. The 50 gene expression datasets for each group were then used as input for consensus analysis that generated two group specific dissimilarity topological overlap maps (dissTOMs) used for module detection by hierarchical clustering.

To compare gene co-expression modules between control and schizophrenia, a module preservation analysis was applied, with control modules as reference and schizophrenia modules as test. Specifically, this method evaluated how well schizophrenia modules were preserved in control modules.

As a preliminary exploratory step, relevant biological mechanisms associated with modules of co-expression in CTRL and schizophrenia groups were summarized for each brain region by gene ontology enrichment analysis with *enrichGO* function implemented in the *clusterProfiler* R package (49).

Details about the WGCNA pipeline are presented in **SM3**.

### Calculation of intra-modular and general connectivity parameters

The next step of gene co-expression network analysis was represented by the calculation of intramodular and general network connectivity parameters and their differences between control and schizophrenia (**Fig.1B, SM4**).

In total, three network metrics were calculated: the first parameter - intramodular degree (intramodular connectivity) - was dependent on modular partition and was normalized by module size (kIn). The two additional parameters were clustering coefficient (C) and total degree (total connectivity) (K). kIn represents the sum of connection strengths between a node (i.e., gene) and all the other nodes in the same module. C is a network parameter of cliquishness measuring how connected the direct neighbors of a node are. K is the sum of connection strengths between a node and all the other nodes in the network (**Fig.1B**).

To better highlight the differences in intramodular connectivity (kIn) between control and schizophrenia, a novel approach for gene module affiliation was applied: control modules were considered as reference and the same gene module affiliation (membership) from control networks was applied to partition the schizophrenia gene co-expression networks (**Fig.1B**). Using this module partition, kIn was calculated from diagnosis specific adjacency matrices with *intramodularConnectivity* function from WGCNA. We also normalized kIn by module size to correct for a bias that shows stronger connectivity in larger modules compared to smaller modules.

To quantify the brain region specific, gene-wise control vs. schizophrenia differences in network parameters (kIn, C and K), a measure of magnitude- the absolute difference- was used to summarize each gene’s differences in network parameters between control and schizophrenia (**Fig.1C**). To explore how consistent the distribution of absolute differences is across the three regions, robust correlations were applied (**SM4** and **Fig.2**).

The most differential nodes, respectively genes outliers for the absolute differences in network parameters, were identified with the *boxplot* function in *base* R (**Fig. 1C**). Region specific patterns of absolute differences’ distributions and outliers for each parameter are visualized in **Fig.3**. The outliers were representative of genes with the most deviant pattern of intramodular or network connectivity in schizophrenia vs. control and were used in subsequent steps to determine their potential biological relevance for schizophrenia.

### Functional profiling of as differentially connected genes (DCG) in schizophrenia co-expression networks

Selection of control vs. schizophrenia DCGs for functional profiling, was based on two criteria: 1. Brain regional consistency (i.e., absolute difference outliers with significant overlap between the three brain regions, denoted as regionally consistent outliers) and 2. Parameter specificity (regionally consistent outliers that are parameter specific by the C, kIn and K network parameters).

1. Regional consistency of absolute difference outliers was tested with the *SuperExactTest* package in R. Routines in this package are used to calculate the exact statistical distributions of multi-set intersections (50) by fold enrichment (FE) (**SM5a**).

2. To select parameter specific regionally consistent outliers, Venn diagram functions (*calculate.overlap* and *VennDiagram*) from the *VennDiagram* package (v.1.7.3) in R were used.

The applied selection pipeline (**Fig.1D**) yielded the final sets of genes with a deviant pattern of network metrics in schizophrenia compared with controls, denoted as differentially connected genes.

Functional profiling of DCG consisted of a. enrichment in schizophrenia genetic signal, b. in cell-type specific markers determined in transcriptomic studies (single-cell RNA-Seq) and c. in gene sets and pathways credibly or previously linked to schizophrenia.

<u>a. post-GWAS analysis.</u> Sets of DCG were tested for association with genetic vulnerability for schizophrenia by two methods- Stratified Linkage Disequilibrium score regression (**LDSC**) (38) and gene-set analysis by Multi-marker Analysis of GenoMic Annotation (**MAGMA**) (39).

Stratified LDSC was used to test the DCG for enrichment in schizophrenia heritability. Following recommendations from the LDSC resource website (https://alkesgroup.broadinstitute.org/LDSCORE), S-LDSC was run for each gene list with the baseline LD model v2.2 that included 97 annotations to control for the LD between variants with other functional annotations in the genome. We defined genomic intervals for each gene within 100 kb upstream or downstream of the gene to capture regulatory regions. We used HapMap Project Phase 3 SNPs as regression SNPs, and 1000 Genomes SNPs of European ancestry samples as reference SNPs, which were all downloaded from the LDSC resource website. For this study, DCG were tested for enrichment in heritability for several complex traits: schizophrenia, the trait of interest and for comparison: ADHD, Bipolar disorder, Major Depressive Disorder, alcohol dependence, epilepsy, Alzheimer’s disorder, autism, intelligence, insomnia, smoking, height, body-mass index (BMI) (details in **SM5b**).

For gene-based association of DCG with schizophrenia genetic risk, linear regression models in MAGMA were used in gene analysis, basic and conditional competitive gene-set analyses (GSA) (38). According to the MAGMA protocol, the first step was to annotate PGC3 schizophrenia GWAS SNPs to genes (excluding the MHC region, respectively genes within chr6:28,477,797-33,448,354), by using as reference the GRCh37 build, and to perform gene analysis. Then, in basic competitive GSA, SNP-wise models were used with default parameters of the program (**SM5b**); finally, in conditional competitive GSA, sets of DCG were tested for enrichment in schizophrenia GWAS signal conditioning on gene sets from synaptic ontologies- SynGO: pre-and postsynaptic processes, and synaptic signaling (41). In addition to network parameter based differentially connected genes (DCG), differentially expressed genes (DEG) between control and schizophrenia in previous transcriptomic studies (15,35) were used in GSA for comparison (**SM5B**).

b. To infer the potential cell-type specificity of DCG two snRNA-Seq resources were used: a transcriptomic study on multiple brain regions, including DLPFC and hippocampus (45) and an atlas focused on glial diversity (46) from hippocampus. From these resources we selected gene markers defined as enriched in DLPFC or hippocampal cell-type subpopulation at the false discovery rate (FDR) < 1e-6 (45) and hippocampus cell-type and subpopulation specific markers for an FDR < 0.05 as described in (46).

### c. Enrichment in schizophrenia-relevant gene sets and pathways

First, relevant gene sets were selected: 1. **PGC3 genes**: 120 prioritized genes from the latest schizophrenia GWAS (42); 2. Gene sets from Transcriptome-wise association studies (**TWAS genes)**: sets of genes with TWAS significance for schizophrenia, shared by the three brain regions (details about TWAS calculation in 15,35); 3. **SynGO genes**: sets of synaptic genes (41) parsed by five synaptic ontologies- metabolism, axo-dendritic transport, pre-synaptic process, post-synaptic process, and synaptic signaling; 4. **Druggable genes**: sets of schizophrenia target genes from the Illuminating Druggable Genome (IDG; 44) database, parsed by the level of development; and 5. Differentially Expressed Genes (**DEGs**) between control and schizophrenia common to the three regions, selected from previous studies from our group (15,35).

Gene set enrichment analysis was performed with routines from *GeneOveralp* package in R that implements a Fisher’s exact test to calculate the overlap between all pairs from two lists of genes (51). The p values and odds ratio were determined in comparison to a genomic background represented by all genes expressed and shared by the three regions (N=18,874). P values were corrected for the number of tests (method: Benjamini-Hochberg False Discovery Rate- FDR).

Biological mapping by gene ontology (GO) analysis: sets of DCG and DEG were compared relative to biological process (GO:BP) enrichment with *compareCluster* function implemented in the *clusterProfiler* R package (49). This function performs enrichment in gene ontologies for multiple gene sets and rank the results by statistical significance (p values) in a comparative manner. The biological theme comparison was visualized with gene-concept network plots and multi-panel dot-plots implemented in the *clusterProfiler* package (**SM5c**).

## DISCUSSION

The molecular architecture of the brain, across regions, cell types and developmental stages varies in a coordinated manner depending on lawful relationships between genes in pathways and networks. Disruptions in these relationships are thought to underlie brain illnesses. In the present study, we aimed to identify genes in the brains of patients with schizophrenia that deviate from the co-expression patterns observed in neurotypical (control) networks. Accordingly, we used three network connectivity parameters to highlight those genes that most significantly differ between controls and schizophrenia and therefore break the rules of connectivity within co-expression networks.

We explored selected differentially connected genes (DCG) from two angles: 1) regional consistency under the assumption that a significant gene overlap across brain regions that differ in cell composition is biologically meaningful and can suggest shared schizophrenia etiopathogenic mechanisms, and respectively 2) parameter specificity under the assumption that network metrics can reveal distinct aspects of the schizophrenia biological substrate. We then looked for DCG association with schizophrenia genetic signal, and enrichment in schizophrenia related ontologies, druggable genome sets and cellular markers.

Our data suggest that schizophrenia differentially connected gene sets by the selected network parameters capture both common and distinct aspects of the biology underlying schizophrenia’s complex disorganization of brain co-expression network architecture. For example, in comparison with C specific DCG and DEG, the kIn specific DCG had the strongest association with schizophrenia GWAS signal: it was the only set significantly enriched in prioritized PGC3 loci genes, for schizophrenia heritability by LDSC and it was the only set associated with schizophrenia genetic signal by MAGMA. Interestingly, this set of differentially connected genes was also enriched for heritability of a variety of neuropsychiatric and neurological disorders, all of which showed previously shared heritability with schizophrenia (22, 52). In contrast, differentially connected genes specific for the clustering coefficient (C specific DCG) showed enrichment only for schizophrenia heritability and also for Alzheimer’s disease heritability. Interestingly, the two DCG sets- kIn and C- were both enriched in heritability of only one normative trait- body-mass index (BMI). The genetic overlap between BMI and schizophrenia was previously documented (53), indicating a potential link between schizophrenia and cardiovascular disorders comorbidity (54).

Biological annotation of differentially connected genes through gene set enrichment analysis and gene ontology analysis brought additional insights about their relations with schizophrenia vulnerability. First, these analyses indicate an interesting segregation by cellular type. While the kIn specific gene set was not relatively enriched for specific cellular markers per se, enrichment in synaptic ontologies suggests a link to neuronal functionality. Conversely, C specific DCG were the only set enriched for cell markers, respectively from the oligodendrocyte lineage, and related oligodendroglia functions. Moreover, gene ontology analysis showed an interesting pathway specialization of kIn and C differentially connected genes. While C specific was biased toward oligodendrocytes differentiation and axon ensheathment, kIn specific was enriched in a mixture of immune processes and modulation of chemical transmission. These varying associations need to be interpreted with some caution as enrichment, per se, is a statistical conclusion that does not establish functional specificity.

Compared with DCG in schizophrenia by co-expression network metrics, differentially expressed genes (DEG) control vs. schizophrenia showed little association with schizophrenia risk in our study, consistent with earlier work (16) and with the assumption that DEG are strongly confounded by drug treatment effects (36). The limited overlap between our DCG and DEG, as well as the overlap between schizophrenia risk genes and DCG, supports the assumption that DCG identified here are potentially less biased by drug effects than DEG.

While a neuronal role in schizophrenia etiopathogenesis has been extensively documented, less is known about the contribution of glial cells, although, recent studies have started to implicate a role played by glia in schizophrenia synaptic dysfunctions (55, 56). Oligodendrocytes, astrocytes and microglia, currently considered key players in shaping synapses structure and modulation of synaptic transmission, can be affected in multiple ways in schizophrenia, starting in neurodevelopment and continuing through adult life (57).

From a network analytical perspective, our study gives a glimpse into the apparent schizophrenia dysconnectivity syndrome at a molecular level, suggesting possible neuronal-glia transcriptional dysregulations. In particular, differences in clustering coefficient between control and schizophrenia across three brain regions seem to be related to the biology of oligodendrocytes. Clustering coefficient is a measure of local density, or “cliquishness”, showing how strongly the neighbors of a node (gene) in the network are connected (58). Since glial cells in general, and particularly oligodendrocytes are credited with a significant role in facilitating synaptic transmission and promoting synaptic plasticity (e.g. through adaptive myelination) (59), it is tempting to speculate that our results based on control vs. schizophrenia differences in clustering coefficient uncover a potential failure of oligodendrocytes to respond to neuronal activity and to support neuronal synapse through adaptive myelination. However, it should be noted that the interplay between neuronal cells and oligodendrocytes in synaptic plasticity is far more complex and involves also astrocytes and microglia (60).

Our study is not without limitations. First, while we extended and innovatively adapted the standard WGCNA pipeline, we measured a reduced number of network parameters. However, considering that little is known about the biological aspects captured by network metrics, our biologically meaningful results are a proof of concept that a more in-depth characterization of gene co-expression networks beyond just finding modules, is worth pursuing in clinical and non-clinical populations. Second, our postmortem brain study is not suitable to account for the dynamics of gene expression regulation and accompanying variations in gene co-expression architecture. Future studies of regulatory dynamics starting with neurodevelopment will be necessary to clarify when the disruptions occur, and how they relate to the etiopathogenesis of schizophrenia. Third, we focused in this study on the most deviant genes by connectivity in schizophrenia, the outliers for absolute difference. However, these may not be the only consequential differences in co-expression network architecture between schizophrenia and control and it is conceivable that testing more cut-offs for filtering the most differentially connected genes will show a more fine-grained image of connectivity patterns altered in schizophrenia.

Fourth, while we focused on regionally shared differentially connected genes, our study also suggested that numerous genes have region specific patterns of deviance in schizophrenia vs. control. These regionally specific differentially connected genes deserve further exploration, considering that differential signatures of gene expression can contribute to the development of structural architecture in the brain and areal functional specialization (34).

Finally, our results derived from bulk RNA-Seq postmortem data could potentially reflect epiphenomenal differences in cellular composition between control and schizophrenia groups. However, recent studies have demonstrated that genetic factors partially drive specific cell-type shifts in the pathogenesis of neuropsychiatric disorders, including schizophrenia (61). This supports our findings of distinct association between parameter-specific differentially connected genes and the genetic signal of schizophrenia, and counters the notion that these network-level differences are merely epiphenomena arising from variations in cell-type composition.

In conclusion, our study of differentially connected genes (DCG) in schizophrenia, brings new insights about possible patterns of dysregulation in this disorder, likely driven by an underlying disruption in the crosstalk between neurons and glial cells that warrants experimental validation.

## Supporting information captions

**S1 text: Supplementary methods (SM).**

**S1 table: Enrichment of differentially connected genes (DCG) in schizophrenia and other complex traits heritability by LDSC.**

**S2 table: Gene set analysis with MAGMA:** a. Gene analysis; b. Basic competitive analysis; c. Conditional competitive analysis.

**S3 table: Results of gene set enrichment analysis (GSEA) of differentially connected genes (DCG).** a. Enrichment of differentially connected genes (DCG) in various gene sets of interest: PGC3 prioritized loci genes, Transcription-wise association genes (TWAS), gene sets from synaptic ontologies (SynGO genes), schizophrenia targets from the Druggable Genome; b. Enrichment of DCG in cell-type markers from DLPFC and hippocampus; c. Enrichment of DCG in glial markers from hippocampus.

**S4 table: Demographics and technical characteristics of postmortem brain samples.**

**S1 figure: General characterization of CTRL and schizophrenia co-expression network modules.** Left panels and upper right panel: cluster dendrograms with schizophrenia modules and CTRL modules matched by color; lower right panel: table with WGCNA results for the region x diagnosis co-expression networks.

**S2 figure:** Summary of GO:BP for control DLPFC modules (scatter plot visualization of biological ontologies with package *rrvgo* (v.1.18.0) in R)

**S3 figure:** Summary of GO:BP for schizophrenia DLPFC modules (scatter plot visualization of biological ontologies with package *rrvgo* (v.1.18.0) in R)

**S4 figure:** Summary of GO:BP for control hippocampus modules (scatter plot visualization of biological ontologies with package *rrvgo* (v.1.18.0) in R)

**S5 figure:** Summary of GO:BP for schizophrenia hippocampus modules (scatter plot visualization of biological ontologies with package *rrvgo* (v.1.18.0) in R)

**S6 figure:** Summary of GO:BP for control caudate modules (scatter plot visualization of biological ontologies with package *rrvgo* (v.1.18.0) in R)

**S7 figure:** Summary of GO:BP for schizophrenia Caudate modules (scatter plot visualization of biological ontologies with package *rrvgo* (v.1.18.0) in R)

**S8 figure:** Module preservation of control modules in schizophrenia gene co-expression networks (Hippo=hippocampus)

**S9 figure:** Correlations between case-control absolute difference for intra-module degree (kIn) across DLPFC, hippocampus and caudate (method Kendall’s tau)

**S10 figure:** Correlations between case-control absolute difference for total connectivity (degree) (K) across DLPFC, hippocampus and caudate (method Kendall’s tau)

**S11 figure:** Correlations between case-control absolute difference for clustering coefficient (C) across DLPFC, hippocampus and caudate (method Kendall’s tau)

