## Supplementary material for "Genes with disrupted connectivity in the architecture of schizophrenia gene co-expression networks highlight atypical neuronal-glial interactions": Suppl_methods: Supplementary_material_01312025.docx

**Supplementary Methods (SM):**

**1. Data selection and processing**

Postmortem brain tissue was collected, dissected, and processed under detailed protocols as described in (1-3).

RNA sequencing: total RNA was extracted from DLPFC gray matter (BA9/46), hippocampus (hippocampus proper and subiculum complex) and caudate (dorsal third, lateral to the lateral ventricle) (1-3). Sequencing libraries were constructed with the TruSeq Stranded Total RNA Library Preparation kit with Ribo-Zero Gold ribosomal RNA depletion.

Raw sequencing reads were quality checked with FastQC (4) and corrected with Trimmomatic (5) if necessary. The quality checked sequencing reads were mapped to the hg38/GRCh38 human reference genome with HISAT2 and Salmon (v2.0.4) (6), and expression for genes was summarized in counts based on GENCODE v25 (GRCh38.p7) (7) and converted to RPKM (Reads Per Kilobase of transcript per Million mapped reads).

Genes with sufficient abundance in more than 80% of samples (RPKM greater than cutoffs determined with *expression_cutoff* function, respectively 0.18 for DLPFC and hippocampus and 0.22 for caudate) were then normalized by log_2_(x+1) transformation. Lastly, samples outliers for gene expression determined by hierarchical clustering of the Euclidean distances measured from the normalized expression data (8), were removed from the analysis.

Genotyping was performed as described in (1). In brief, it comprised genotype imputation with IMPUTE2 and Shape-IT after removing low quality and rare variants. Common variants with MAF>5% present in >90% of samples, and in Hardy-Weinberg Equilibrium (HWE p value >0.001) were retained. Multidimensional scaling (MDS) was performed to calculate ancestry components (genomic PCs).

**2. Data “cleaning” (removal of unwanted variance explained by biological and technical covariates)**

Linear regression models with normalized expression data as dependent variable and covariates of interest (diagnosis) and non-interest (biological and technical) were used to “clean” (adjust) the normalized gene expression from the three brain regions. Before applying the cleaning step, a multicollinearity test implemented in R *car* package (*vif* function) was performed. After discarding variables with VIF ≥10, several covariates of non-interest were retained for expression adjustment: 1. Age, sex, 10 ancestry components (genomic PCs), technical covariates (RIN, mitoRate) and 11 quality surrogate variables (qSVs) for DLPFC and caudate; 2. Age, sex, 10 ancestry components, 12 qSVs and technical covariates (RIN, totalAssignedGene and mitoRate) for hippocampus.

qSVs were calculated with Quality Surrogate Variable Analysis (QSVA) that consists of identifying transcript features most susceptible to RNA degradation by estimating a “degradation matrix” quantified as the coverage of the susceptible features in customized sequencing libraries and performing a principal component analysis on the “degradation matrix” that yields a number of k principal components named “quality surrogate variables” (qSVs). Quality surrogate variables (qSVs) were extracted from region-specific RNA degradation matrices calculated as described in (9).

The number of qSVs was pre-determined with the *num.sv* function that uses the method of Buja and Eyuboglu (9).

After specifying the linear models for expression adjustment, *cleaningY* function (jaffelab R package v.0.99.30) (10) was used to preserve effects from covariates of interest and remove unwanted variance explained by covariates of non-interest. The procedure described by Jaffe et al (2017) (9) was based on the observation that biological signal of interest could correlate with artifacts and therefore would be removed during the step of data cleaning if not preserved.

The general equation of linear models for cleaning

***Expression*** *~ β_0_ +* ***β1*Diagnosis*** *+ β_2_*Age + β_3_*Sex + β_4_*RIN + β_5_*totalAssignedGene + β_6_*mitoRate + Σβ_i_*snpPC_i_+ Σβ_j_*qSV_j_*

was adapted to include the retained covariates for each region, while preserving the intercept and diagnosis (parameter P=2 in *cleaningY* function).

**3. Gene co-expression network analysis with WGCNA**

For the co-expression network analysis, we used WGCNA package version 1.69 (11-12). Adjusted expression data were used as input for network construction.

After calculating the beta power, CTRL and SCZ datasets were bootstrapped by using a sampling procedure with replacement (*sample.int* function in *base* R); 50 CTRL and 50 SCZ gene expression bootstraps were then used for a robust WGCNA, following a procedure adapted from (13).

For each group, a consensus network analysis from the 50 gene expression bootstraps was performed with *consensusTOM* function (correlation type = bi-weight midcorrelation; type of network = signed, β powers: 10 for DLPFC and caudate, 14 for hippocampus, selected with soft thresholding to correspond to an R^2≥0.8; networkCalibration = "single quantile", consensusQuantile = 0.5). The results were represented by two consensus networks (one for CTRL, one for SCZ) for each brain region. Consensus networks were then used to calculate topological overlap matrix (TOM) dissimilarity (dissTOM). Finally, dissTOMs were used to detect modules by hierarchical clustering with a deep split parameter=3 and a minimum module size of 50 nodes (gene). The modules were labeled with pre-specified colors implemented in WGCNA.

To compare CTRL vs. SCZ modules for each region, *modulePreservation* function in WGCNA was used. With this function, network-based, pair-wise module preservation statistics were calculated by taking as input CTRL gene expression and modules as reference set, and SCZ gene expression and modules as test set (parameters for module preservation: network type = signed, correlation type = bi-weight midcorrelation, number of permutations = 100). Most relevant output from the network preservation statistics is represented by measures of preservation for density (i.e. if a module remains densely connected in the test network) and connectivity (i.e. if the hub gene status is preserved between reference and test networks, where hub genes are the most connected genes in a module) (14). Four density and four connectivity preservation statistics were calculated in the module preservation analysis. Individual Z scores from these computations were summarized in a composite measure called Z_summary_. By convention, Z_summary_ indicates no preservation for a range between 0-2, weak preservation between 2-10 and strong preservation >10 (14). Another composite statistic (importantly, not dependent on module size), is the median rank.

Results of the module preservation were displayed on plots with module size on x axis and preservation Z summary, respectively preservation median rank on y axis.

Functional enrichment of CTRL and SCZ modules for each brain region was performed with the *enrichGO* function from the Bioconductor *clusterProfiler* package v.4.8.3 (15). This function calculates the gene ontology overrepresentation (i.e., biological process- GO-BP) with a hypergeometric test by which functional terms are tested for statistically significant enrichment in gene lists of interest while using as statistical domain the region-specific background (the total gene set: DLPFC = 22,542; caudate = 19,954, hippocampus = 20,708). False Discovery Rate (FDR) correction was used for multiple comparisons correction. Parameters used for *enrichGO* function were minimum gene set to be analyzed- 100, respectively maximum gene set- 5,000; method for p value adjustment- Benjamini-Hochberg; p value cutoff=0.001, q-value cut-off=0.001. Visualization of enrichment results was performed with functions from the *rrvgo* R package v.1.12.2 (16).

**4. Calculation of intra-modular and general network parameters**

To calculate intra-modular degree and total degree, the *intramodularConnectivity* WGCNA function was used. This function takes as input adjacency matrices (correlation matrices raised at beta power) and nodes’ module affiliation to calculate the intramodular degree by summing for each node (gene) adjacencies entries to the other genes in the module. The output of this function is represented by intramodular connectivity (degree), total degree (total connectivity), extra-modular connectivity and the difference between intra- and extra-modular connectivity. Specifically for our study, intramodular degree was scaled (divided) by module size, which represented Km connectivity parameter.

Clustering coefficient was calculated with *clusterCoef* function in WGCNA, which also uses as input adjacency matrices.

The case-control absolute differences in network metrics was subsequently calculated for each gene as described in the main manuscript and checked for consistency across the three brain by using Kendall tau correlations. The correlation analyses were performed and results visualized with functions from *PerformanceAnalytics* (v.2.0.8) and *GGally* (v.2.2.1) R packages that calculate and visualize robust correlations. The reason for selecting robust (i.e., non-parametric) correlations was dictated by the non-normal shape of distribution for the selected absolute differences.

Outliers for case-control absolute differences in connectivity parameters were considered the most differentially connected genes (DCG) between CTRL and SCZ networks.

**5. Functional profiling of DCG in SCZ co-expression networks**

a. Brain regional specificity. Function *supertest* from the *SuperExactTest* package (v.1.1.0) (17) was used to compute the exact statistical distribution of multi-set intersections, where the sets were represented by outliers for absolute difference in network metrics from the three brain regions- DLPFC, hippocampus and caudate. Essentially, this function implements algorithms to calculate expected and observed overlap size, the fold enrichment (FE) and test statistics for all 2^n^-1 possible intersections for n sets (i.e., in our case n=3). The assumption for common outlier genes shared by the three regions was that outliers for absolute difference in network metrics were independently and randomly sampled from a total of 23,256 genes representing all common genes across the 3 regions plus all region-specific genes.

b. post-GWAS analysis. Stratified LDSC was performed as recommended in LDSC tutorials (<https://github.com/bulik/ldsc/wiki/Partitioned-Heritability>) (18). First, genome positions for gene sets were converted from hg38 to hg19 reference assembly with *biomaRt* (v.2.62.0), to be concordant with the GWAS summary statistics. Then genomic regions for differentially connected genes (DCG) (gene location plus 100kB around start and end of the transcription site) were annotated chromosome-wise to the 1000 Genomes (EUR ancestry). Stratified LDSC was applied to estimate the contribution of DCG annotations to trait heritability conditional on the 97 annotations in the baseline model.

MAGMA software (v.1.10) (19) was used for gene-set enrichment analysis with sets of **DCG** in comparison with genes from synaptic ontologies- SynGO (20) grouped in synaptic signaling, pre- and postsynaptic processes and axo-dendritic transport and differentially expressed genes (CTRL compared with SCZ) in postmortem RNA-Seq transcriptomic studies. First, all genes expressed in the three brain regions were aligned to the GRCh37 (hg19) assembly and used for annotation of SNPs from PGC3 SCZ GWAS, (i.e., all SNPs in a window around the transcription region of the gene: +1 kB/ +0.5 kB), as recommended in MAGMA protocol; then, a first step included gene analysis of all expressed genes (N=18090) with the SNP-wise model; the association of each gene with SCZ genetic vulnerability by considering the LD structure inferred from the European reference panel from the 1000 Genomes, was performed with a Bonferroni correction α=0.05/18090; the results from the gene-based analysis were used in a basic competitive gene-set analysis, with gene sets from **DCG** (C specific, Km specific), and differentially expressed genes shared by the 3 regions (**DEG**) and SynGO pathways (synaptic signaling, axo-dendritic transport, pre- and postsynaptic processes). A Bonferroni correction α=0.05/6 was used. Finally, using the same gene sets, a competitive conditional gene-analysis was performed, using the SynGO processes as conditions.

c. Gene Ontology analysis:

Functional enrichment of DCG was performed with the *clusterProfiler* package v.4.8.3 (15). For this part, *compareCluster* function was used*.* This function compares gene sets enrichment in functional terms (i.e., GO:BP) by using the hypergeometric test.

**Selective references for supplementary methods:**

1. Jaffe AE, Straub RE, Shin JH, Tao R, Gao Y, Collado-Torres L, et al (2018): Developmental and genetic regulation of the human cortex transcriptome illuminate schizophrenia pathogenesis. Nat Neurosci 21:1117-1125.
2. Collado-Torres L, Burke EE, Peterson A, Shin J, Straub RE, Rajpurohit A, et al (2019): Regional Heterogeneity in Gene Expression, Regulation, and Coherence in the Frontal Cortex and Hippocampus across Development and Schizophrenia. Neuron 103:203-216.e8.
3. Benjamin KJM, Chen Q, Jaffe AE, Stolz JM, Collado-Torres L, Huuki-Myers LA, et al (2022): Analysis of the caudate nucleus transcriptome in individuals with schizophrenia highlights effects of antipsychotics and new risk genes. *Nat Neurosci* 25:1559-1568
4. Babraham Bioinformatics (2016). FastQC (Babraham Institute). <https://www.bioinformatics.babraham.ac.uk/projects/fastqc/>.
5. Bolger AM, Lohse M, Usadel B (2014): Trimmomatic: a flexible trimmer for Illumina sequence data. Bioinformatics 30:2114-2120
6. Kim D, Paggi JM, Park C, Bennett C, Salzberg SL (2019): Graph-based genome alignment and genotyping with HISAT2 and HISAT-genotype. Nat Biotechnol 37:907-915.
7. Frankish A, Diekhans M, Ferreira AM, Johnson R, Jungreis I, Loveland J, et al (2019): GENCODE reference annotation for the human and mouse genomes. Nucleic Acids Res 47:D766-D773.
8. Horvath, S (2011). Chapter 1, pp.1-34, in Weighted Network Analysis. Applications in Systems Biology. Springer: New York, Dordrecht, Heidelberg, London.
9. Jaffe AE, Tao R, Norris AL, Kealhofer M, Nellore A, Shin JH, et al (2017): qSVA framework for RNA quality correction in differential expression analysis. Proc Natl Acad Sci U S A 114:7130-7135.
10. jaffelab package: <https://github.com/LieberInstitute/jaffelab>
11. https://horvath.genetics.ucla.edu/html/CoexpressionNetwork/Rpackages/WGCNA/Tutorials/Consensus-NetworkConstruction-man.pdf
12. WGCNA: <https://cran.r-project.org/web/packages/WGCNA/WGCNA.pdf>
13. Parikshak NN, Luo R, Zhang A, Won H, Lowe JK, Chandran V, et al (2013): Integrative functional genomic analyses implicate specific molecular pathways and circuits in autism. *Cell* 155:1008–1021.
14. Langfelder P, Luo R, Oldham MC, Horvath S (2011): Is my network module preserved and reproducible? *PLoS Comput Biol* 7:e1001057.
15. Yu G, Wang LG, Han Y, He QY (2012): clusterProfiler: an R package for comparing biological themes among gene clusters. OMICS 16:284-287.
16. Sayols, S (2023). rrvgo: a Bioconductor package for interpreting lists of Gene Ontology terms. microPublication Biology. [10.17912/micropub.biology.000811](https://doi.org/10.17912/micropub.biology.000811).
17. Wang M, Zhao Y, Zhang B (2015): Efficient Test and Visualization of Multi-Set Intersections. *Sci Rep* 5:16923.
18. Bulik-Sullivan BK, Loh P-R, Finucane HK, Ripke S, Yang J, Consortium SWG of the PG *et al.* LD Score regression distinguishes confounding from polygenicity in genome-wide association studies. 2015; **47**: 291–295.
19. Finucane HK, Reshef YA, Anttila V, Slowikowski K, Gusev A, Byrnes A, et al (2018): Heritability enrichment of specifically expressed genes identifies disease-relevant tissues and cell types. *Nat Genet* 50:621-629.
20. Koopmans F, van Nierop P, Andres-Alonso M, Byrnes A, Cijsouw T, Coba MP, et al (2019): SynGO: An Evidence-Based, Expert-Curated Knowledge Base for the Synapse. *Neuron* 103:217-234.e4.
