## Supplementary material for "Genes with disrupted connectivity in the architecture of schizophrenia gene co-expression networks highlight atypical neuronal-glial interactions": Figures_suppl: Figures_suppl_01312025.pptx

### Slide 1
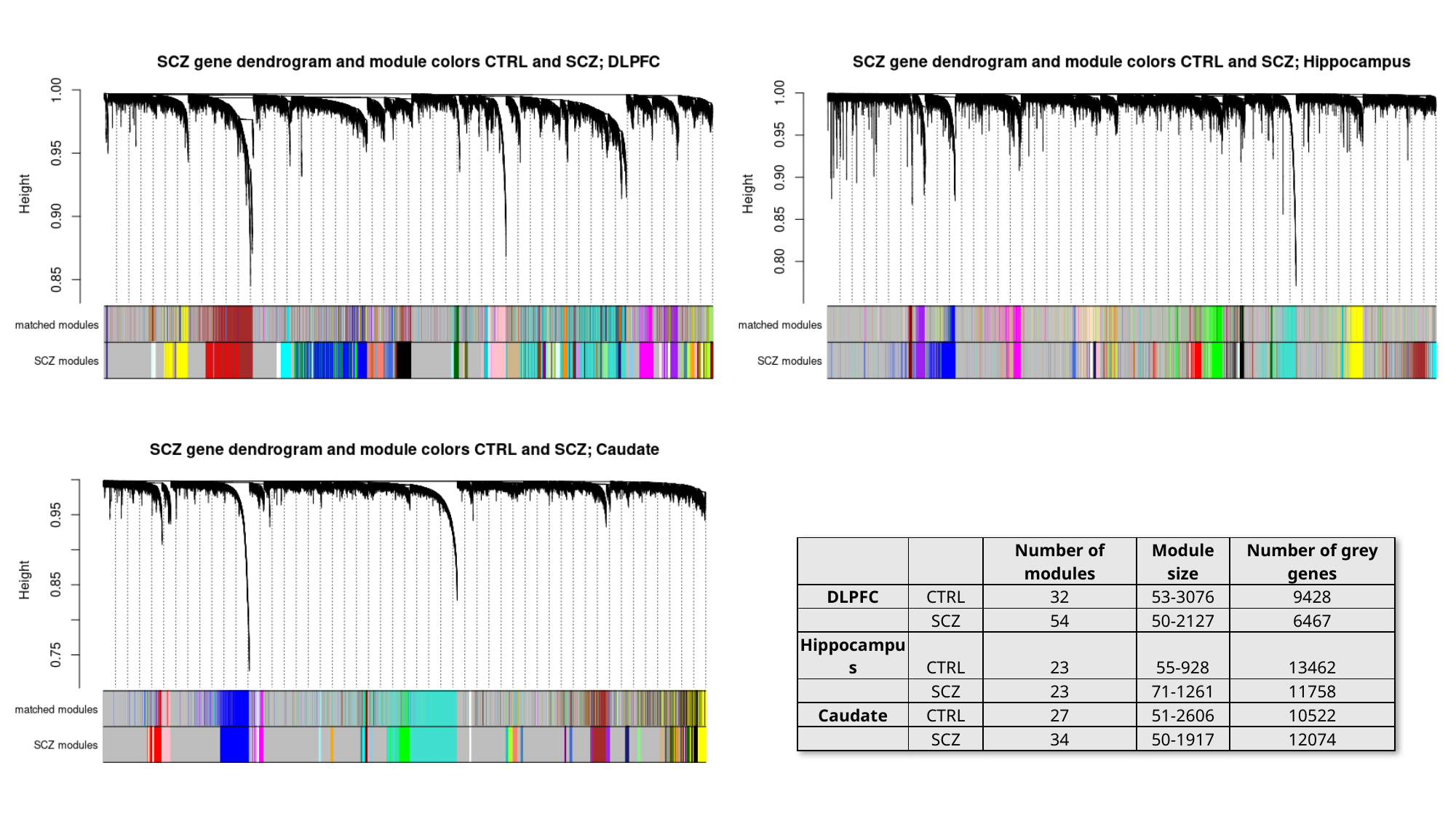

| | | Number of modules | Module size | Number of grey genes |
| --- | --- | --- | --- | --- |
| DLPFC | CTRL | 32 | 53-3076 | 9428 |
| | SCZ | 54 | 50-2127 | 6467 |
| Hippocampus | CTRL | 23 | 55-928 | 13462 |
| | SCZ | 23 | 71-1261 | 11758 |
| Caudate | CTRL | 27 | 51-2606 | 10522 |
| | SCZ | 34 | 50-1917 | 12074 |

### Slide 2
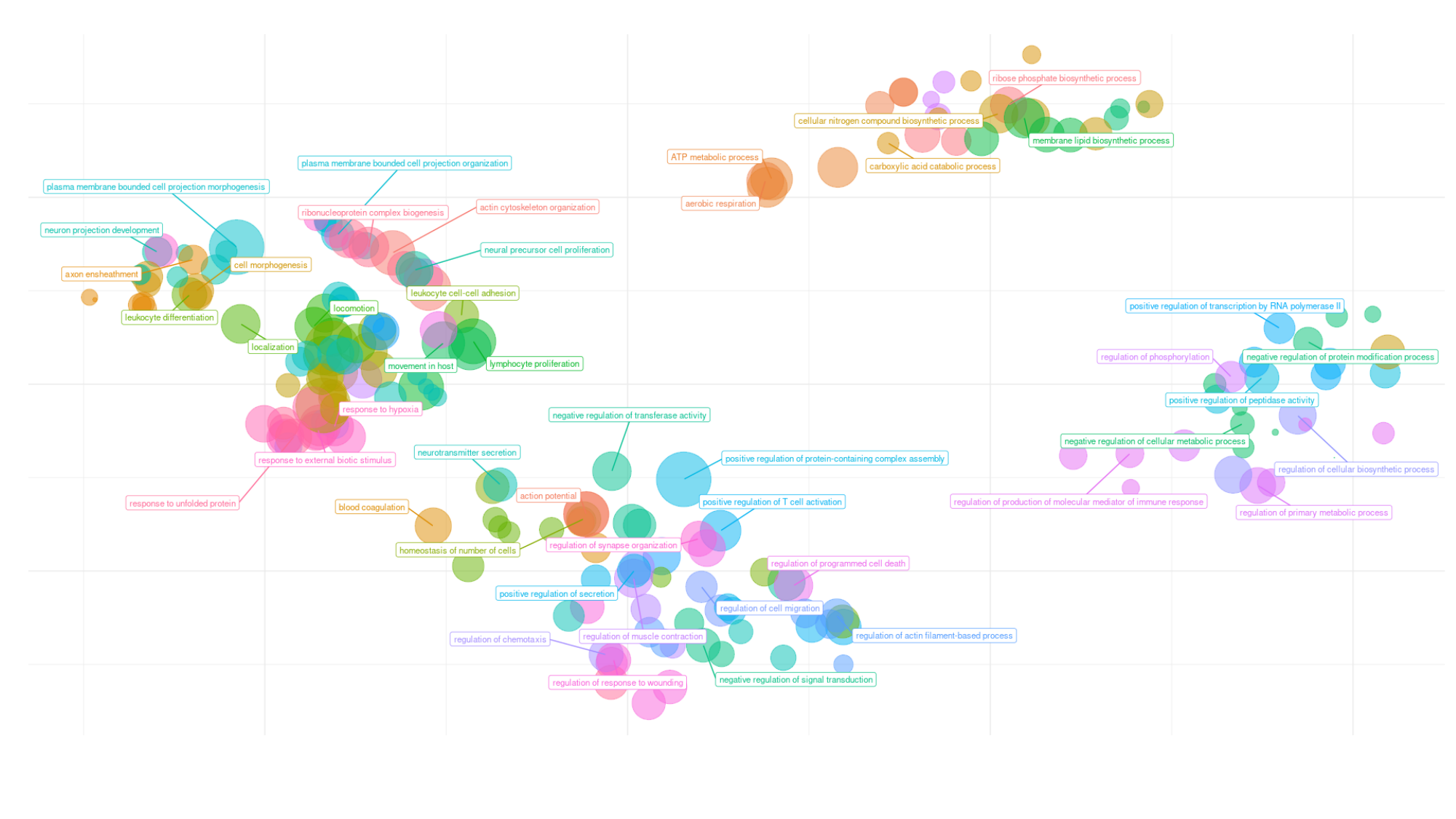

### Slide 3
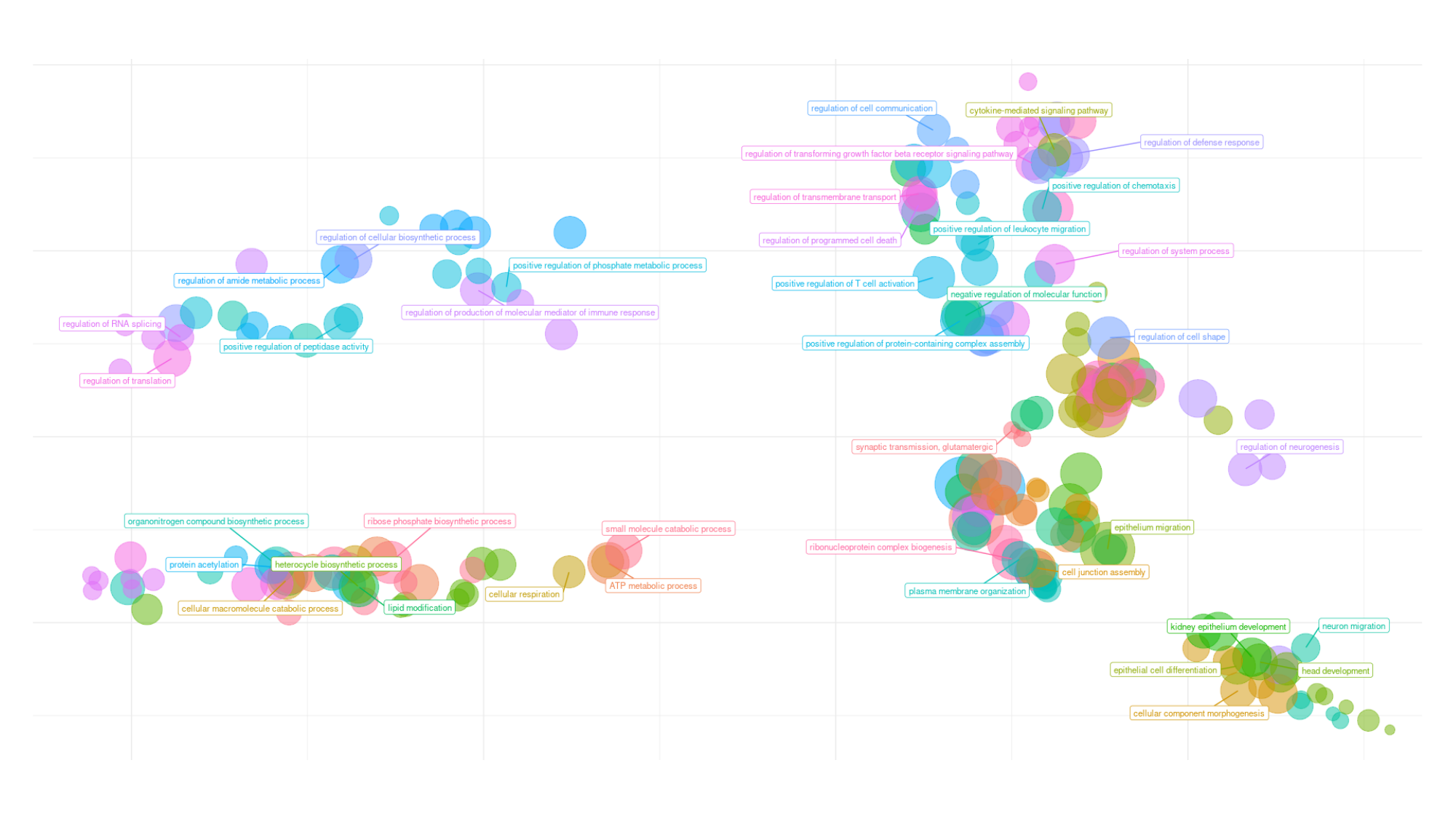

### Slide 4
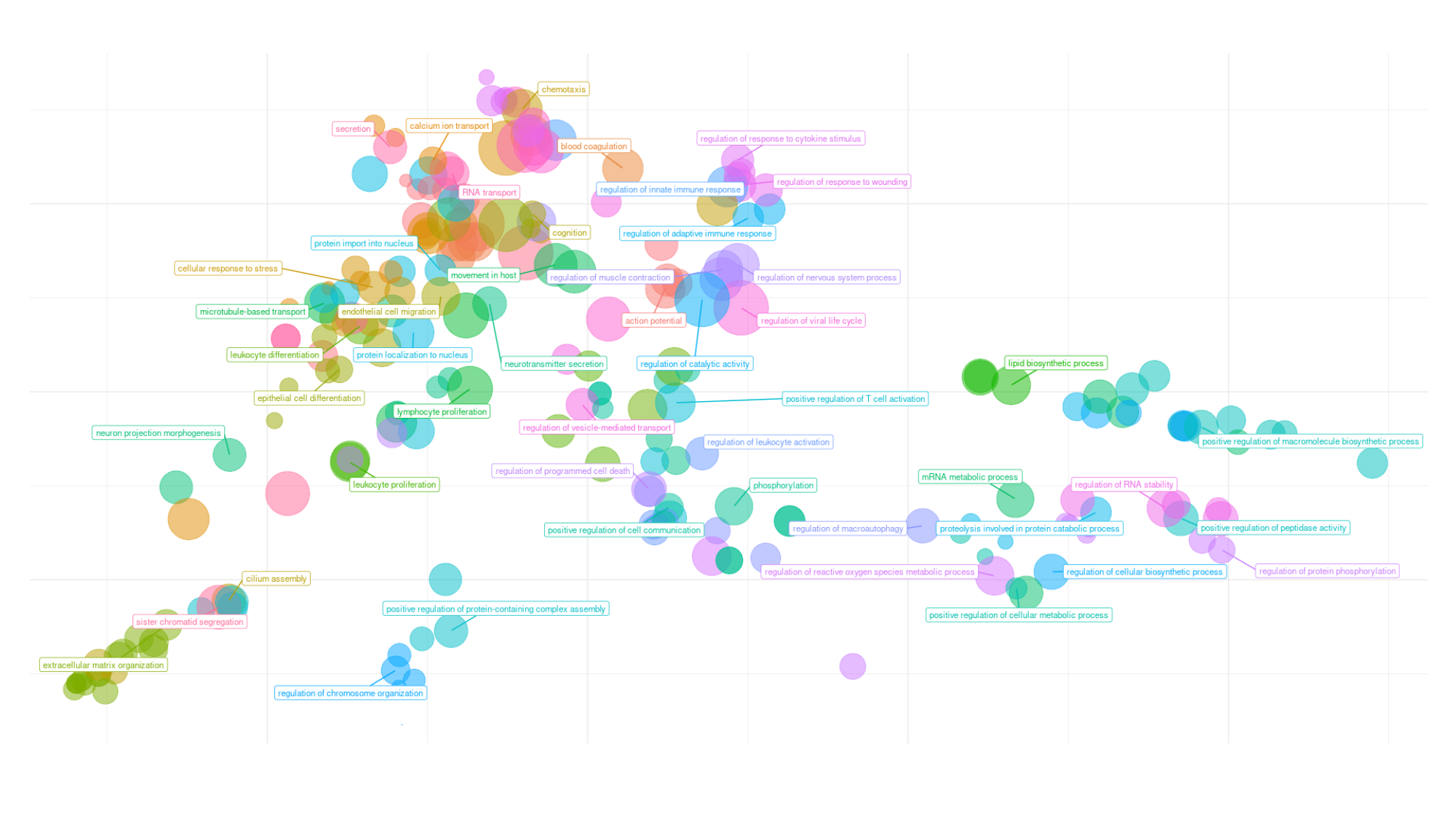

### Slide 5
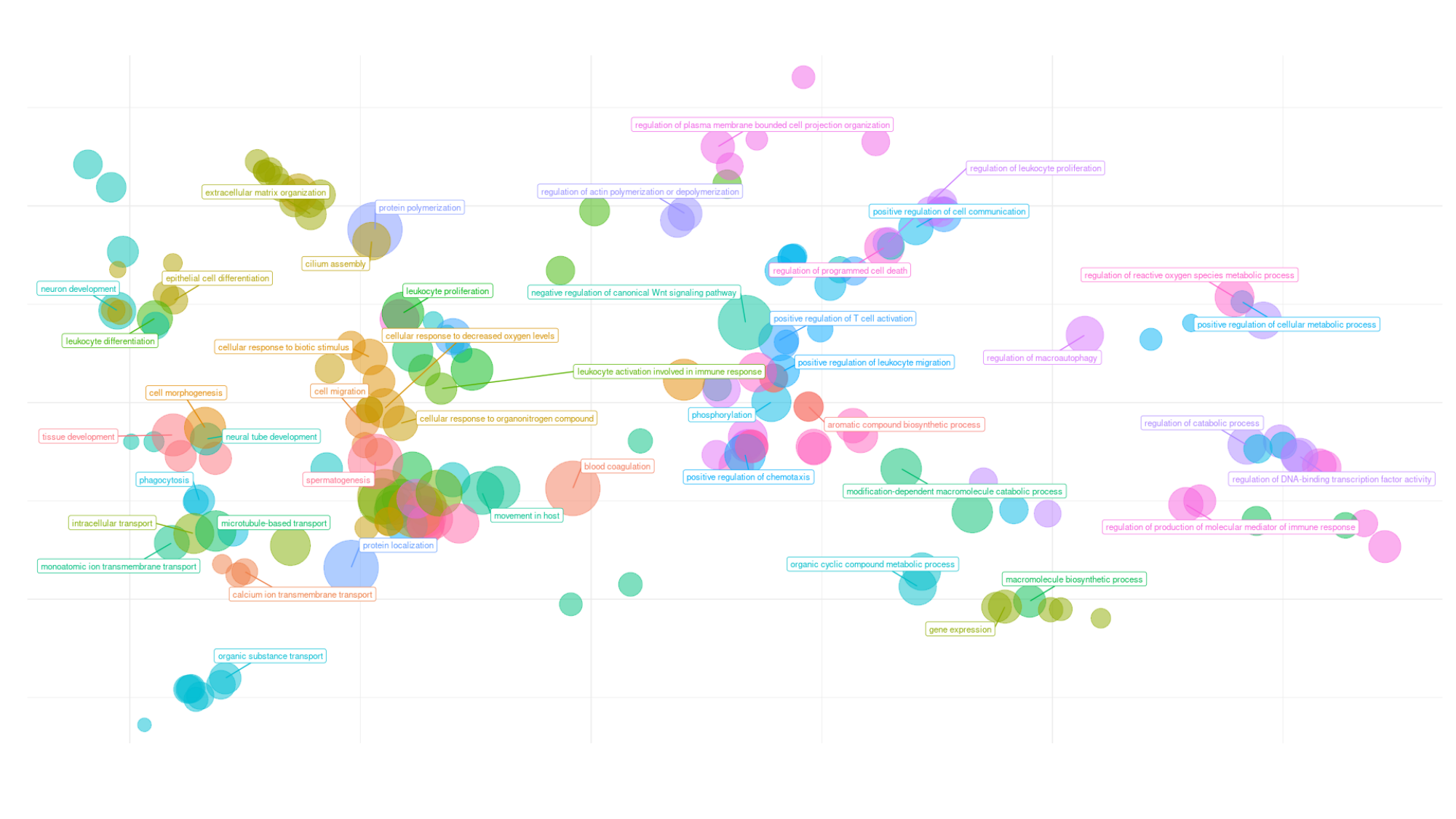

### Slide 6
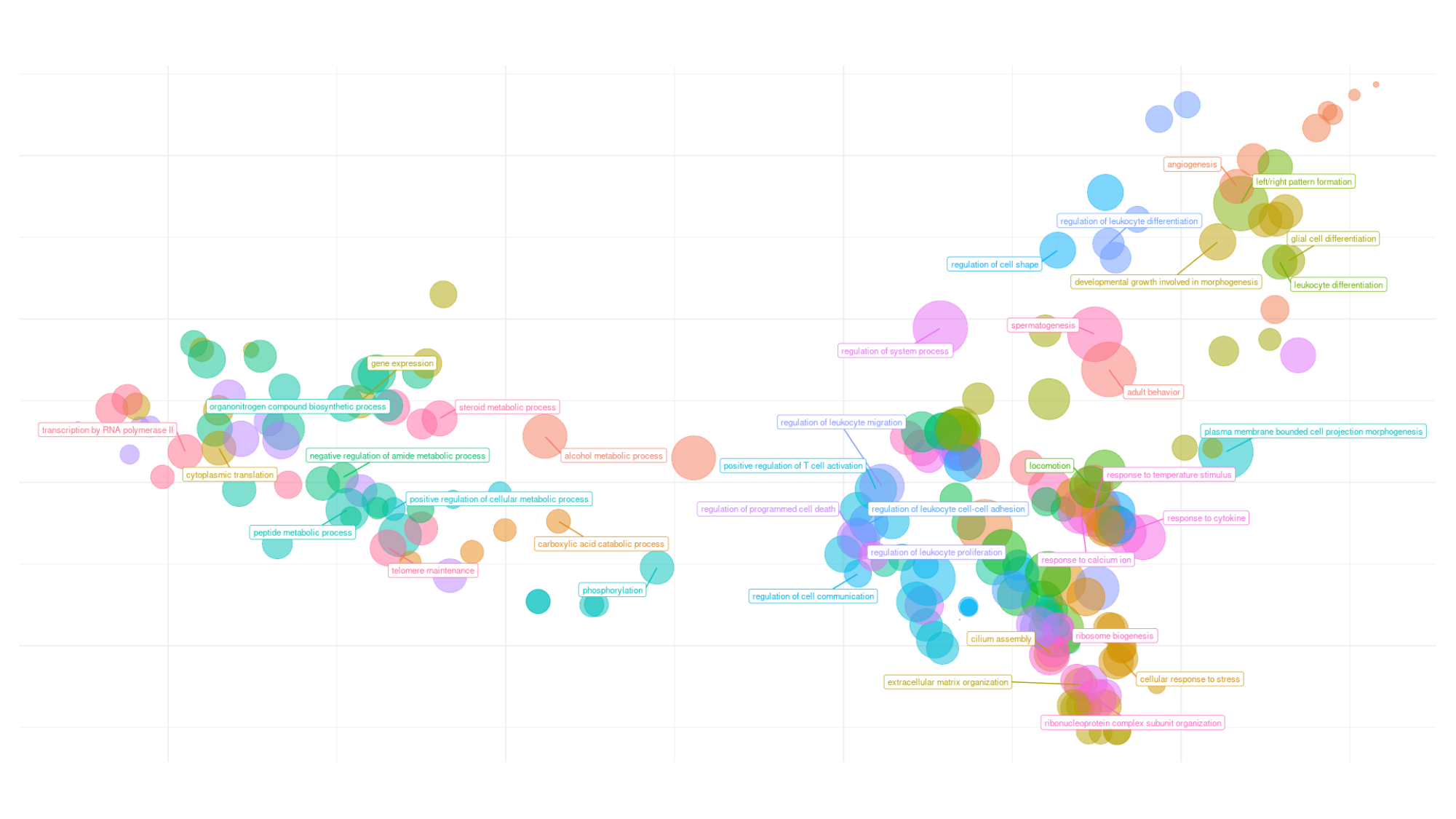

### Slide 7
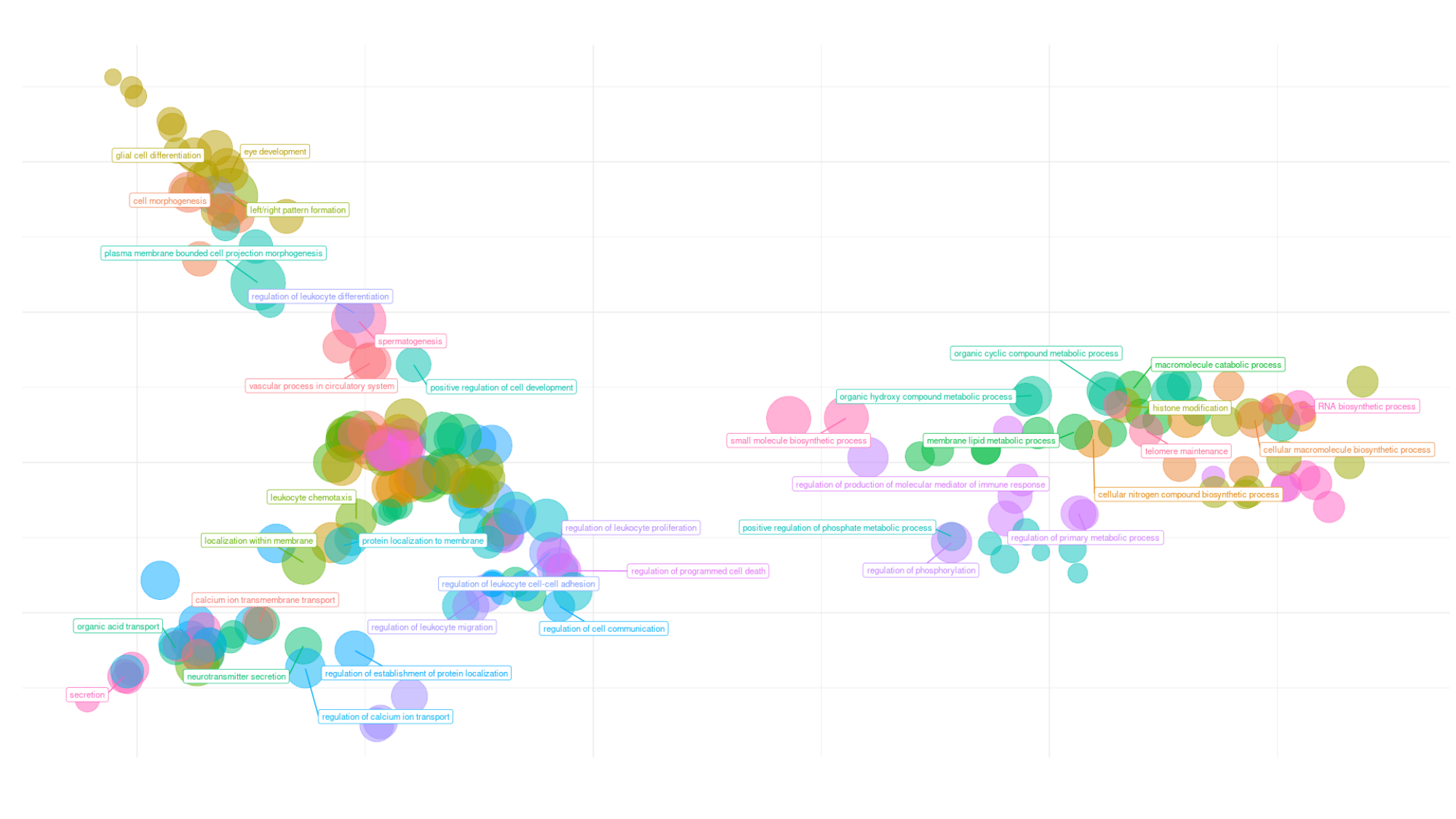

### Slide 8
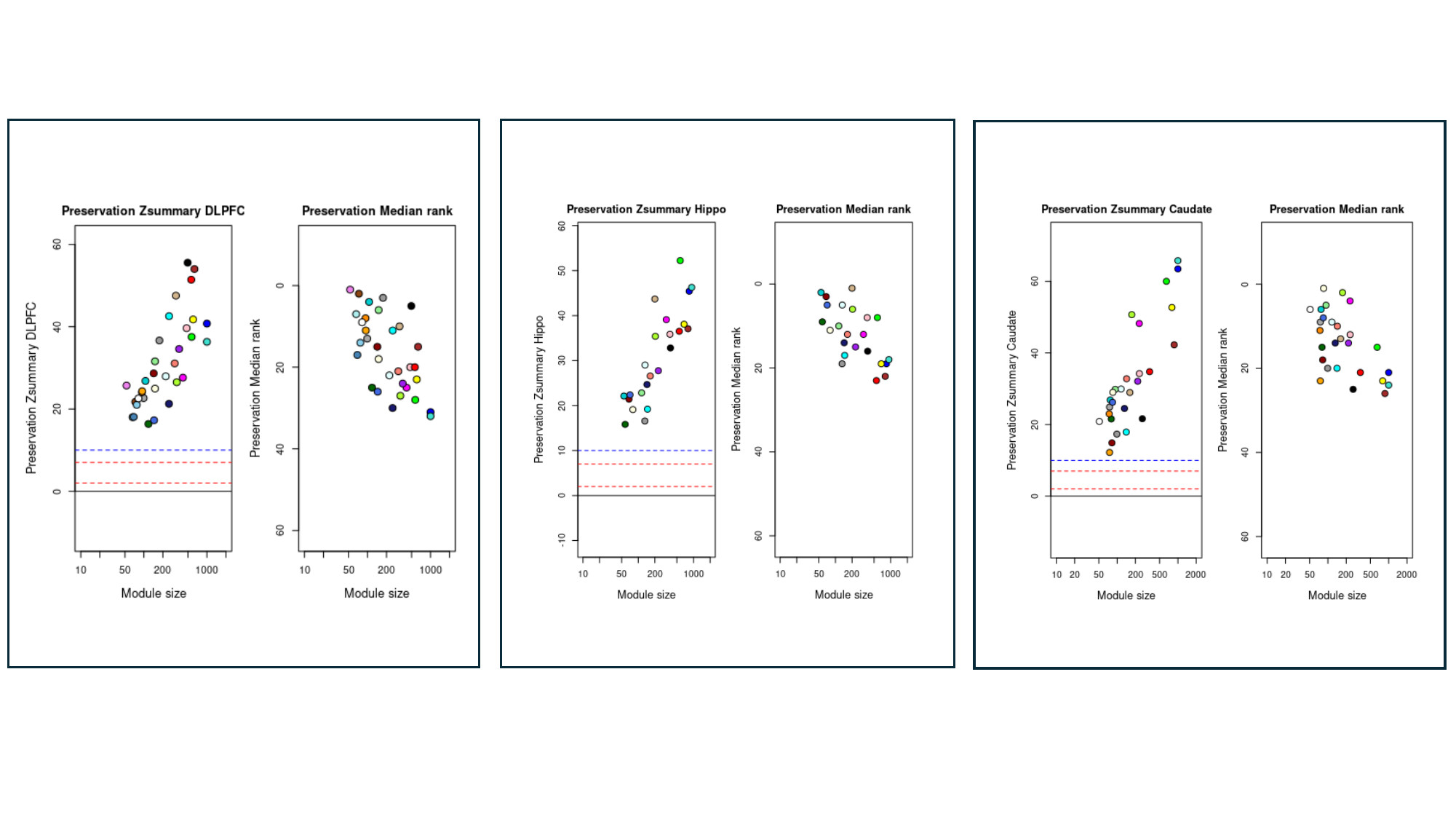

### Slide 9
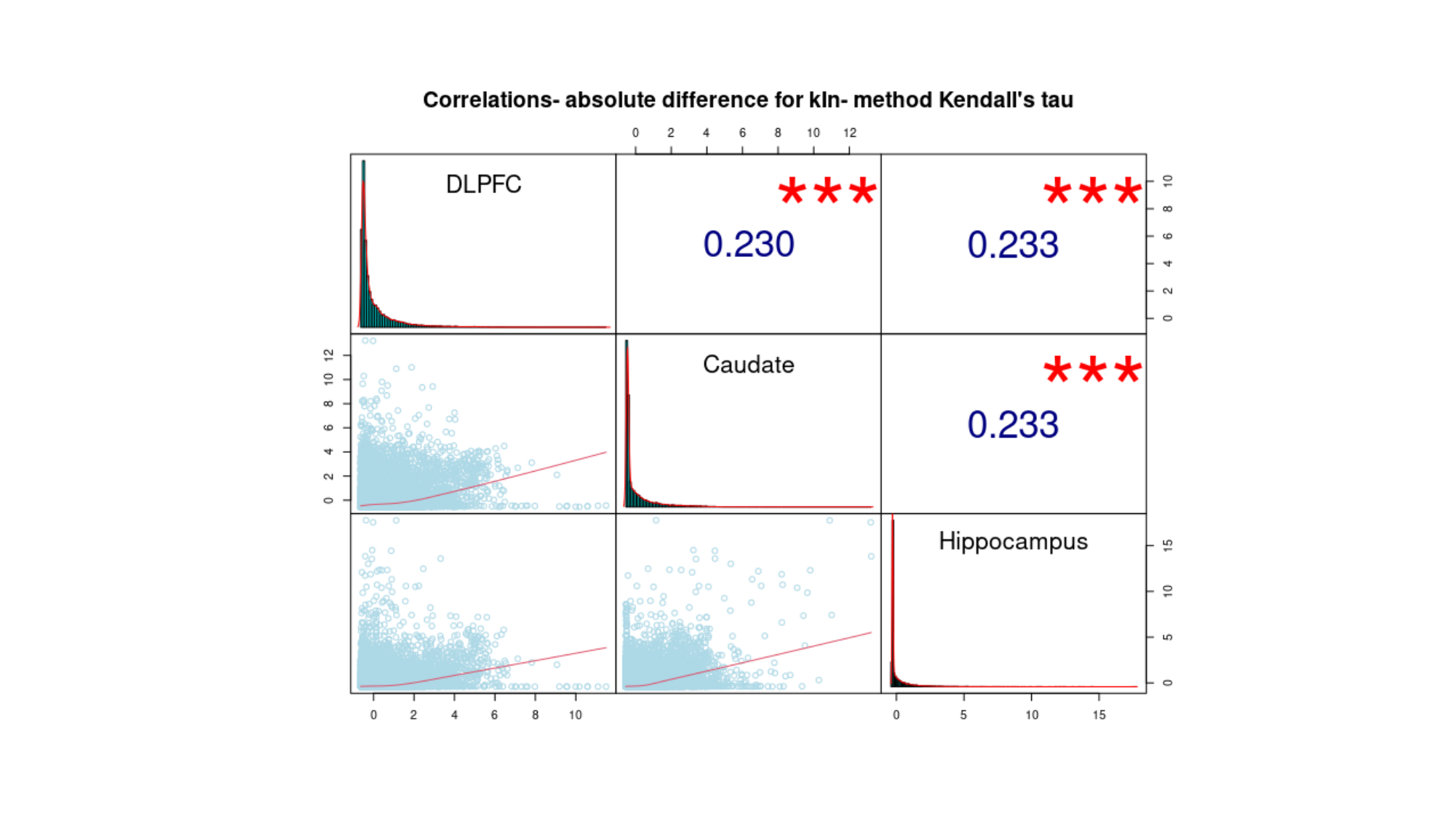

### Slide 10
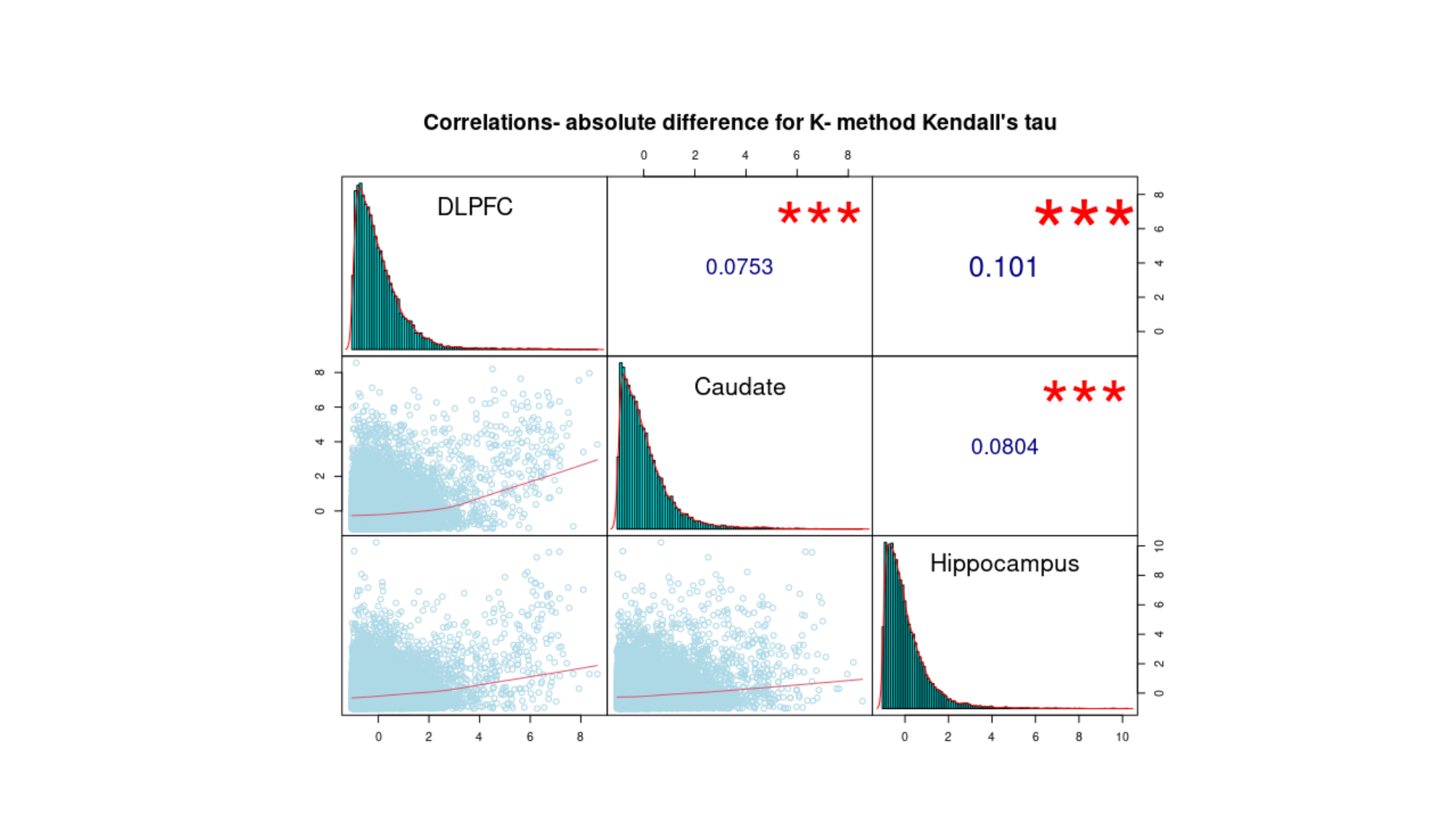

### Slide 11
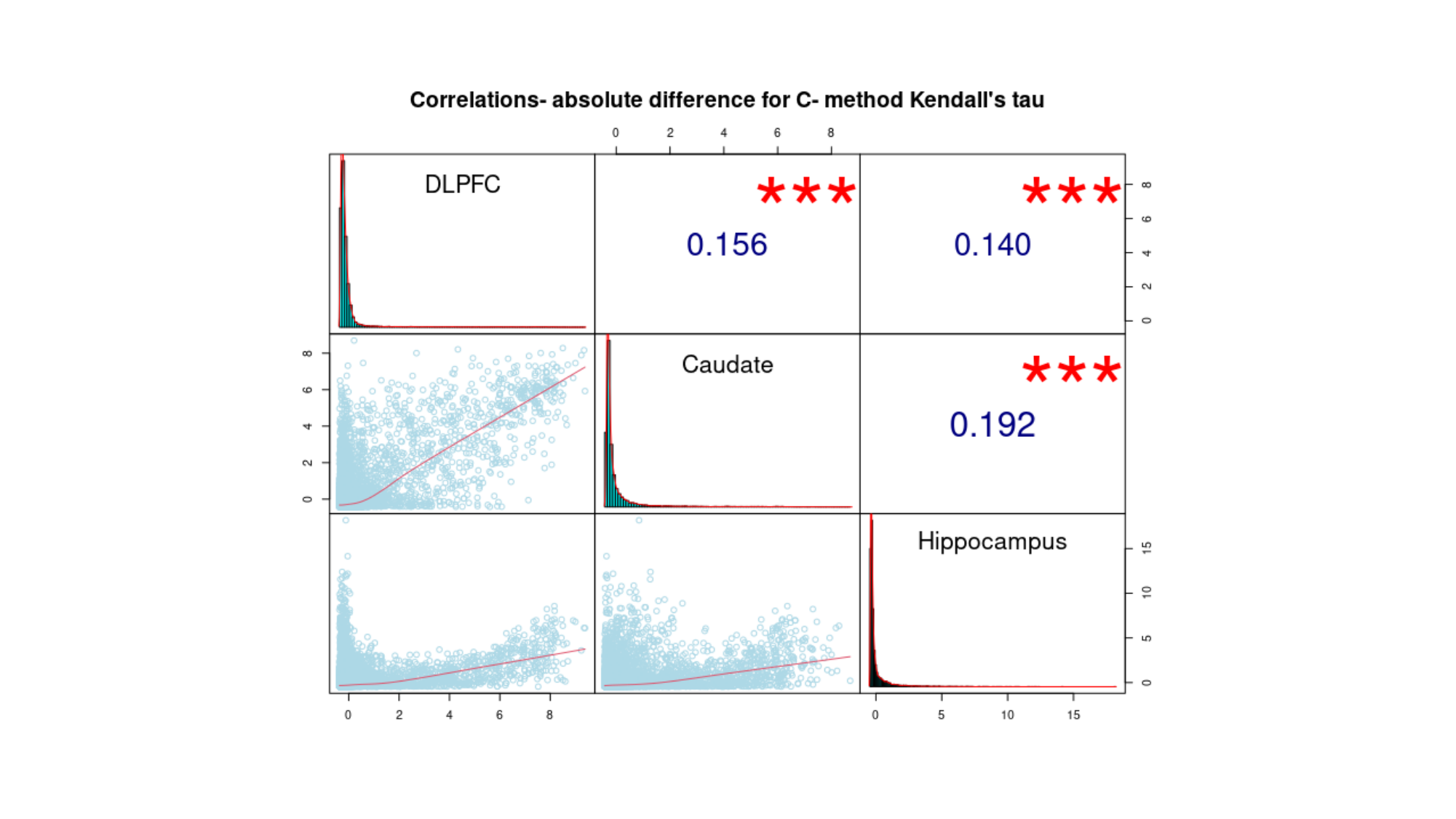
